# The gut microbiota and sleep in infants: a focus on diurnal rhythmicity patterns

**DOI:** 10.1101/2025.07.11.664337

**Authors:** Fannie Kerff, Christophe Mühlematter, Anja Adamov, Deborah Fast, Serafina Plüss, Petra Zimmermann, Salome Kurth, Nicholas A. Bokulich

## Abstract

Emerging evidence supports a bidirectional relationship between the gut microbiome and sleep, partly mediated by the microbiota-gut-brain axis. Circadian rhythms appear to influence early developmental processes and may play a role in shaping this relationship. While infancy is a critical window for both gut microbiota establishment and the maturation of sleep regulation, the role of gut bacteria in modulating infant sleep remains poorly understood.

In this longitudinal study, we performed continuous within-subject sampling (monitoring across 48 hours) in 20 infants at 2, 4, and 6 months of age to examine associations between the establishment of gut microbiota and sleep patterns. We hypothesized that microbial diversity and sleep rhythmicity would be linked across both short (diurnal) and long (monthly) time scales. Gut microbiota profiles were characterized using 16S rRNA gene sequencing; gut melatonin concentrations were measured; and sleep metrics were quantified through wearable actimetry, parent-reported 24-hour sleep diaries, and the Brief Infant Sleep Questionnaire (BISQ). Parents additionally completed the Baby Care Questionnaire (BCQ), assessing parenting style, and the Ages and Stages Questionnaire (ASQ), evaluating infant behavioral development.

In some infants, alpha diversity followed diurnal rhythmic patterns. Although bacterial rhythmicity was not significantly associated with sleep rhythmicity, as quantified with the Circadian Function Index, infants with higher alpha diversity had more robust sleep rhythmicity (β_Faith PD_= 0.0721, SE = 0.0337). Infant age emerged as the strongest predictor of both gut microbiota diversity (observed features: β_age_ = 0.101, SE = 0.0172) and gut melatonin concentrations (β_age_ = 0.638, SE = 0.162). For the same age, gut microbiota temporal volatility – an indicator for bacterial community instability – was not associated with sleep patterns. This study demonstrates promising short– and long-term links between gut microbiota diversity and sleep-wake rhythm maturation in infancy. Further research is needed to elucidate the mechanistic role of the gut microbiome in sleep development.

## Introduction

Microbial colonization of the gastrointestinal tract begins at birth and is influenced by multiple factors including maternal microbiota, gestational age, mode of delivery, feeding practices (breastfeeding vs. formula), and antibiotic exposure (Bokulich et al., 2016; Fahur Bottino et al., 2025; La Rosa et al., 2014; Zimmermann & Curtis, 2018). During the first 1,000 days of life, the gut microbiome undergoes its most rapid period of development, playing a critical role in immune, endocrine, and metabolic functions, and other host developmental processes (Robertson et al., 2019).

Growing evidence suggests that the gut microbiome is associated with both sleep (Sen et al., 2021) and cognitive abilities (Ahrens et al., 2024). Notably, the first year of life – a period of development and stabilization of the gut microbiome – coincides with important milestones in sleep regulation and neurodevelopment (Meltzer et al., 2021; Schoch et al., 2020). In particular, brain oscillations during sleep actively regulate neuroplasticity, thereby supporting learning and healthy neurodevelopment (Timofeev et al., 2020). The bidirectional microbiome-sleep relationship is partially mediated by the microbiota-gut-brain axis and is shaped by environmental and lifestyle factors (Sen et al., 2021).

The development of sleep regulation primarily includes a gradual establishment of rhythm, including for example increased consolidation of sleep and wake bouts, increasing preference towards nighttime sleep and less fragmented nighttime sleep (all along a continuum). Several pathways of gut microbiota-host interaction suggest possible influence on sleep development. For instance, in infants, human milk oligosaccharides (HMOs) promote the growth of *Bifidobacteria* (Lawson et al., 2020), which produce short-chain fatty acids (SCFAs) such as propionate (Wiciński et al., 2020). Fecal propionate levels have been positively associated with sleep duration in infants (Heath et al., 2020), supporting the relationship between microbial metabolites and early-life sleep regulation. Similarly, infants receiving a prebiotic blend of indigestible oligosaccharides in infant formula have shown faster consolidation of daytime waking periods (Colombo et al., 2021) and potential beneficial effects on sleep duration (Nocerino et al., 2020), suggesting beneficial roles of *Bifidobacteria* and SCFAs production, although further clarification on probiotics is needed (Papagaroufalis et al., 2014; Sung et al., 2014). Recent findings also indicate that infants’ sleep-wake patterns are associated with regularity of feeding schedules (Mühlematter et al., 2023). Since certain gut bacterial taxa show diurnal oscillations (Mühlematter et al., 2025), and several SCFAs fluctuate rhythmically throughout the day (Kaczmarek et al., 2017), it is likely that the systemic circadian rhythm plays a role in the molecular mechanisms underlying sleep–microbiome interactions.

While systemic melatonin is recognized for its crucial role in regulating circadian rhythm, research on intestinal melatonin production by and/or associations with gut bacteria is only emerging (Al-Andoli et al., 2025; Zimmermann et al., 2024). Additionally, small amounts of melatonin are present in certain foods (e.g., pistachios, cherries, grapes), which may also contribute to intestinal and systemic melatonin levels (Meng et al., 2017). Even though the pineal gland, producing melatonin, and melatonin receptors are present at birth, the ability of the gland to release and produce melatonin rhythmically is just developing across the first year of life (Attanasio et al., 1986; Kennaway et al., 1996). Using a cross-sectional approach, it was proposed that fecal melatonin increases with age (high inter-individual variability), with higher levels earlier in the day (diurnal variations), and higher levels with more time passed since the last bowel movement (accumulation of melatonin) (Al-Andoli et al., 2025). Still, contrasting evidence on diurnal fluctuations were reported, and the specific pathways and bacterial species involved in melatonin metabolism (both production and degradation) in the gut remain unclear (Zimmermann et al., 2024).

Despite evidence that gut microbiota diversity is associated with sleep in adults (Gao et al., 2022; Grosicki et al., 2020; Liu et al., 2019; Smith et al., 2019) and school-aged children (Deng et al., 2022; Hua et al., 2020; Lawrence et al., 2022; Valentini et al., 2020; Wang et al., 2022; Xiang et al., 2023), very few studies have examined microbiota-sleep interactions in infancy (0-1 year). Among these, longitudinal studies in infants have shown links between gut bacterial diversity – representing microbial profile maturation – and the maturation of behavioral sleep patterns, including nighttime preference for sleep, but also neurophysiological markers of sleep homeostasis (slow wave activity) (Schoch et al., 2022). Moreover, emerging evidence suggests that diurnal rhythmicity patterns already develop in the gut microbiota and are impacted by the diet (Heppner et al., 2024) and intertwined with the host’s circadian rhythm (Mühlematter et al., 2025).

Altogether, despite infancy being a critical period for both microbiome development and sleep regulation establishment, the role of gut bacteria in modulating infant sleep remains underexplored. Notably, high-frequency sampling is needed to capture time series and potential rhythmicity across the 24-hour day, and only longitudinal trajectories adequately capture the rapid and multi-dimensional transitions in the infant gut microbiota and sleep. To address this gap, this study aimed to explore (1) microbial rhythmicity and (2) temporal volatility patterns (bacterial community variability over time) along age, then (3) quantify the effect of feeding and sleep history on the microbiota, and finally (4) determine the link between gut microbiota patterns and sleep in a longitudinal cohort of infants. We hypothesize that across this early-life developmental period, the gut microbiome and sleep rhythm establishment are linked on a short-term (diurnal) time scale, and that infants with a less diverse or more volatile gut bacterial profile exhibit a delayed establishment of sleep maturation, captured by the longitudinal design across several months. With the aim of advancing microbiome-based therapies to improve sleep, this study focuses on how the gut microbiome influences sleep, rather than vice versa.

## Methods

### Study design

We launched an observational longitudinal cohort study to investigate the synchronization of gut microbiota and sleep in infancy. From May 2023 to April 2024, this study followed 20 healthy term-born infants across development with assessments scheduled at 2, 4, and 6 months of age. Inclusion criteria required infants to be generally healthy, vaginally delivered, and primarily breast-fed at the time of enrollment.

All procedures were conducted in accordance with the Declaration of Helsinki and approved by the responsible cantonal ethics committee (BASEC 2019-02250). Written parental consent was obtained after explanation of the study procedures and before effective enrollment. Families received non-monetary gifts to maintain adherence across assessments.

### Experimental design

Gut microbiota was evaluated through biological sampling (stool); infant sleep was assessed with objective (actimetry via wearable devices) and subjective (24-hour diaries and surveys) parameters. A graphic study design is displayed in Fig. 1**A**.

**Fig. 1.**
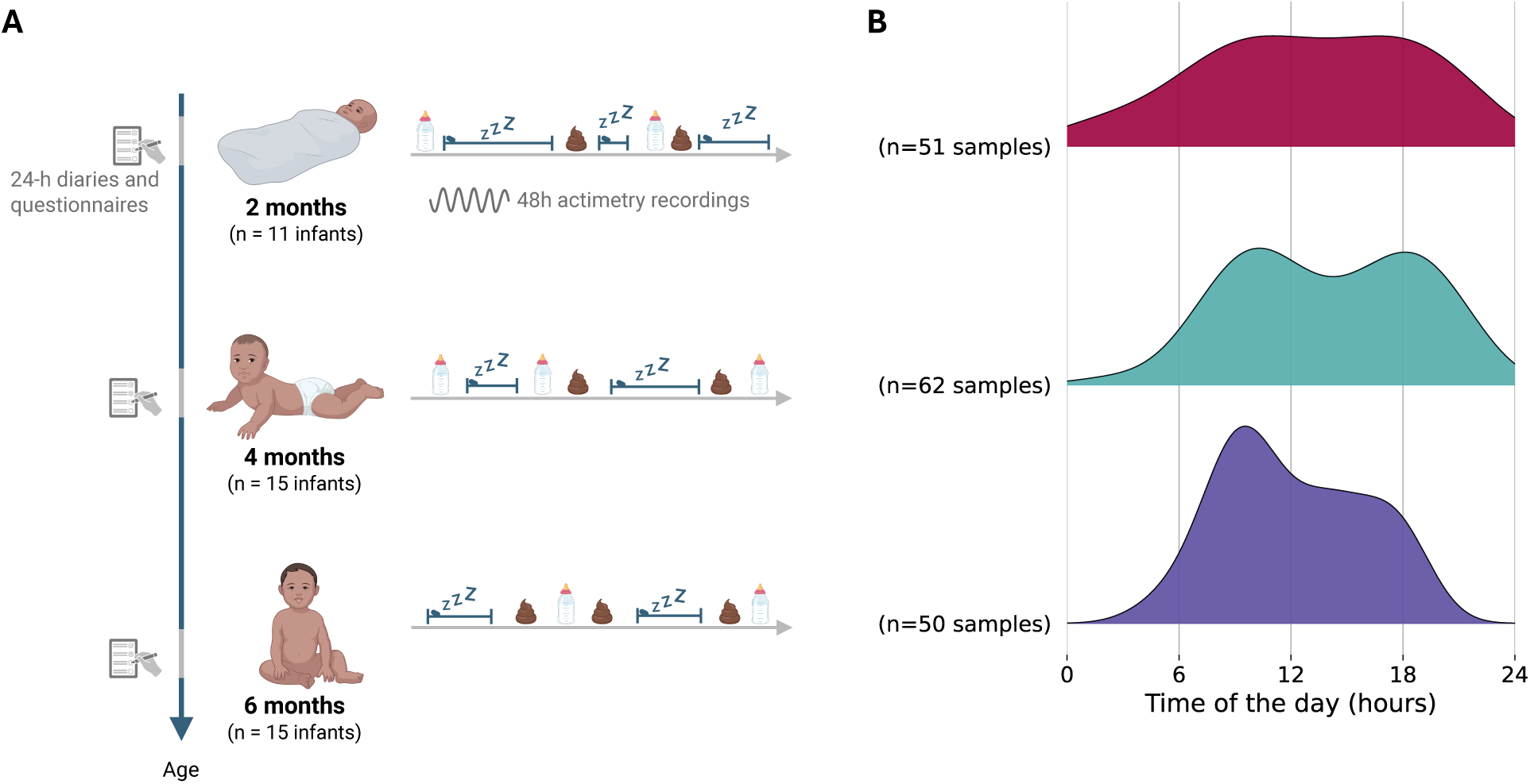
Study overview. **A** Study design. **B** Stool sample collection density according to clock times across age assessments (i.e., ages 2 months, 4 months, and 6 months).

### Stool sample collection

Parents collected stool samples from the infant’s diaper at all ages using sterile disposable pipettes or laboratory spatulas (Schoch et al., 2022). Samples were kept in 5 mL Eppendorf tubes, placed in Whirlpak bags and temporarily stored in the families’ fridge, until transported to the laboratory (University of Fribourg) within 76 hours in cooling boxes for temperature maintenance. Aliquots were generated and stored at –80°C.

### Sleep data collection

At each age assessment, continuous sleep data were collected across several days using ankle actimetry (GENEactiv accelerometer, Activinsights Ltd, Kimbolton, UK 43 x 40 x 13 mm, Micro-Electro-Mechanical Systems sensor, 16 g, 30 Hz frequency) combined with a 24-hour sleep-wake diary which was adapted from Werner et al. (2008). Parents documented actimeter removal in the diary, which included detailed 15-min interval reporting of sleep or wake periods, feeding times, crying, and bedtimes. Using data from both actimetry and the 24-h diary, sleep and wake periods and feeding times were identified following our laboratory’s established standards (Mühlematter et al., 2023; Schoch et al., 2019). Additionally, the Brief Infant Sleep Questionnaire (BISQ), a clinically-validated parent-reported questionnaire capturing infants/toddler (0-29 months) habitual sleep was used at all ages (Sadeh, 2004).

### Additional questionnaires and feeding diary

Demographic information was provided by parents in an online survey at each assessment, which was generated using SoSci Survey. Further, at all ages, parents filled a parenting style questionnaire (BCQ) (Winstanley & Gattis, 2013), as well as standardized questionnaires on behavioral development (ASQ) (Squires et al., 1995). Feeding times (clock times of infant meals) were reported by the parents in the sleep diary at all ages.

## Data processing

### Infant gut microbiota

#### Stool processing: DNA extraction and 16S rRNA gene sequencing

DNA was extracted from approx. 80 mg (60 to 100 mg accepted; 80 µL if sample was liquid) of stool using MagMAX™ Microbiome Ultra Nucleic Acid Isolation Kit with bead plates following manufacturer’s instructions with one modification: bead beating 2x 5 minutes at 30 Hz. 400 µL of lysate were transferred to the KingFisher Deep Well Plates and stored at – 20°C. DNA extraction was done on the KingFisher Apex (binding, washing, elution). Eluted DNA was collected in KingFisher standard plates and stored at 4°C (fridge) overnight.

We used the HighALPS ultra-high throughput library preparation protocol (Flörl et al., 2024) to profile bacterial communities via sequencing the V4 region of the 16S rRNA gene using 515F and 806R primers (Apprill et al., 2015; Parada et al., 2016). DNA from each sample was amplified using barcoded forward and reverse primers targeting the 16S rRNA gene (1-step PCR; 34 cycles). PCR products were cleaned with 0.7x AMPure XP beads on the KingFisher Apex, quantified with the Qubit assay, combined into one final pool after serial dilutions, and manually cleaned with 0.7x beads, followed by an elution in 100 µL nuclease-free water. Qubit (dsDNA HS Assay) and TapeStation (HS D1000) analyses confirmed sufficient concentration and quality of the final pool, which was submitted for library pooling.

#### Sequences processing: gut microbiota composition and diversity

Sequence data were analyzed using QIIME 2 (Bolyen et al., 2019). Sequences were demultiplexed, denoised (truncation at 150 bp using the q2-dada2 plugin (Callahan et al., 2016)), and rarefied at a sequencing depth of 3,035 reads. Features with a frequency less than 10 across all samples were discarded. Taxonomic classification was performed using a naive Bayes classifier (Bokulich et al., 2018) trained on the V4 region extracted from SILVA version 138.1 (Pruesse et al., 2007; Quast et al., 2013) (using RESCRIPt (Robeson et al., 2021)), and sequences not assigned to a phylum were removed. Finally, diversity metrics were computed using the q2-boots plugin (Raspet et al., 2024), averaging the metrics over 100 computations. Calculated alpha diversity metrics included observed features, Shannon entropy, Pielou evenness and Faith phylogenetic diversity; beta diversity metrics measured were Bray-Curtis dissimilarity (Sørensen et al., 1948), Jaccard similarity index (Jaccard, 1908), unweighted UniFrac (C. Lozupone & Knight, 2005) and weighted UniFrac (C. A. Lozupone et al., 2007).

#### Gut microbiota rhythmicity

Gut microbiota rhythm per infant and age was assessed through a cosine fitting of bacterial alpha diversity to detect 24-hour oscillation, following the equation:

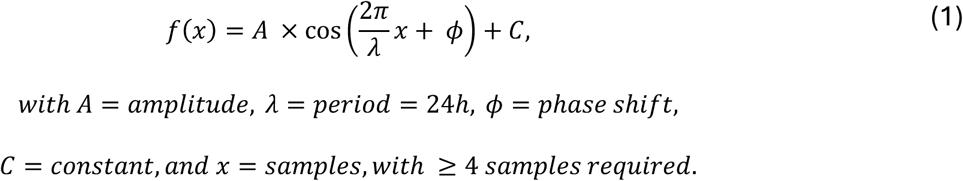

Bacterial diurnal rhythmicity was quantified with R^2^ values computed for each cosine fit. High R^2^ values represented a good cosine fit of alpha diversity over time.

#### Gut microbiota temporal volatility

For each infant, we quantified the variability of the gut bacterial community over time (48h) at each age. To do this, the centroid of each infant was defined for each beta diversity metric at each age. The centroid was computed as the mean position of the three first principal components of those samples. The gut bacterial variability of each infant at each age was estimated by measuring the median distance of the samples to their corresponding centroid (i.e., the centroid of the (≥ 2) samples of that infant at that age). This measure of variability in the microbiota of an infant (across 48h) is henceforth referred to as gut bacterial “temporal volatility” of the infant, at that age.

### Gut melatonin

Samples with enough remaining stool were analyzed for melatonin. Gut melatonin concentration was determined using a radioimmunoassay (RIA) RK-MEL2, manufactured by NovoLytiX GmbH (Witterswil, Switzerland) following the manufacturer’s instructions (IFU, version 2022-03-07) with specific adaptations for stool samples. For each sample, 50 mg of stool was homogenized in 1 mL of RK-MEL2 Incubation Buffer; the mixture was vigorously vortexed for 10 seconds and left for 10 minutes at room temperature to maximize homogenization. The latter step was repeated 3 times, and the homogenate was centrifuged at 13,000 rpm (7,000x g) for 10 minutes in an Eppendorf Minifuge; the same centrifugation procedure was done to the supernatant. Finally, the supernatant was used for RIA, with a correction factor of 0.77 (as established by standards (Markovic et al., 2024)) to adjust for matrix effects, validated through spiking recovery experiments.

### Infant sleep

#### Sleep rhythmicity: Circadian Function Index (CFI) score

Based on actimetry data, the Circadian Function Index (CFI) was computed, which measures the regularity of activity-rest patterns with a score from 0 to 1 (Ortiz-Tudela et al., 2010), and was used as a proxy for sleep-wake rhythm maturation for each infant at each age (Mühlematter et al., 2025). The CFI was quantified for infants who were assessed during at least three continuous days at an age. The CFI is defined as:

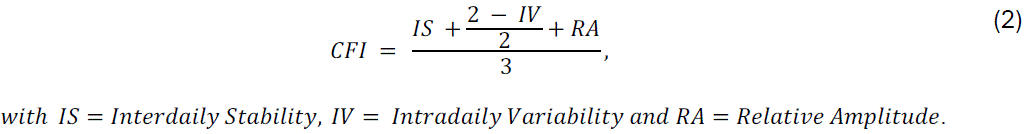

Interdaily Stability (IS) measures how consistent activity patterns are across days; higher IS indicates more regular, stable daily rhythms. The Intradaily Variability (IV) indicates how often transitions occur between rest and activity within a day; higher IV means more fragmented patterns; lower IV reflects more consolidated periods. Relative Amplitude (RA) reflects the difference between peak activity and rest; higher RA shows a clearer distinction between active and rest periods. The CFI is henceforth referred to as “sleep rhythmicity”.

#### Infant sleep quality: babySQUID

An infant Sleep QUality InDex (“babySQUID”) was determined for each age as the composite score of four variables derived from the BISQ sleep questionnaire: nighttime sleep duration, sleep onset latency, bedtime and number of nighttime awakenings. The babySQUID estimated sleep quality over one week. Infants with maximally two missing infant sleep variable(s) were still included, with the missing values replaced by the median of that variable. All variables were min-max scaled, then all variables except nighttime sleep duration were reverse-coded (sleep onset latency, bedtime and number of nighttime awakenings), and the normalized values were averaged with equal weights to produce a single score from 0 to 1, reflecting overall sleep behavior. In this context, we hypothesized that earlier bedtimes indicated a more consolidated sleep, as previously linked to stronger neuronal connectivity in infants (Schoch et al., 2024).

### Other variables

We used parent-reported diaries to determine the infant’s sleep and feeding history prior to each stool sample. To estimate the sleep history, we compared the stool sample clock time with the parent-rated entries (nighttime sleep and awakenings, wake up time, and daytime naps) to assess the time spent awake prior to each sampling. Then, we used the related diary information on falling asleep times and naps to estimate the duration of that last sleep. The same methodology was used to estimate the feeding history (time since the last feeding) using the parent-rated feeding clock times in the diaries. Finally, feeding rhythmicity was estimated by calculating the standard deviation of the intervals between feedings for each infant for each day, at each age assessment.

The parenting style (attuned *versus* structured) was estimated using the attunement score of the BCQ questionnaire. Attunement represents reliance on infant cues and close physical contact; parents with a more attuned caring/parenting style would be more prone to change their behavior based on the needs of the infant. Infant behavioral developmental stage was estimated using the composite score (sum) of the five sub-scores of the ASQ questionnaire.

### Statistical analysis

For all analyses, missing values in exposure or outcome variables led to the exclusion of that sample/age for that analysis, while missing variables in the covariates were averaged by the median value of that variable. Note that each infant’s gut microbiota diversity at a given age was estimated using the median of their alpha diversity values at that age. Statistical significance was determined at the 5% level.

### Diurnal patterns along age

To investigate the links between diurnal rhythmicity patterns in the gut microbiota and infant sleep rhythmicity, a Linear Mixed-Effects Model (LMM) was created adjusting for the infants’ repeated measures, feeding rhythmicity, parenting style (BCQ score), age and sex, following the equation:

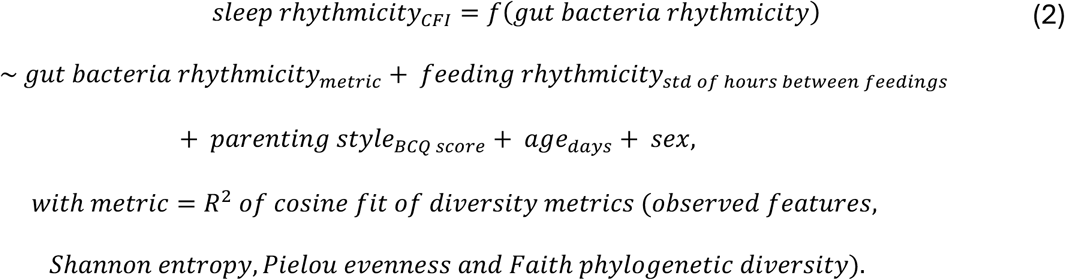

Effects of sleep and feeding history on the gut microbiota

To identify which directly preceding sleep and feeding history factors predominantly determine the infant gut microbiota, LMMs for diversity metrics were created, adjusting for the infants’ repeated measures, as well as age and sex, following the equation:

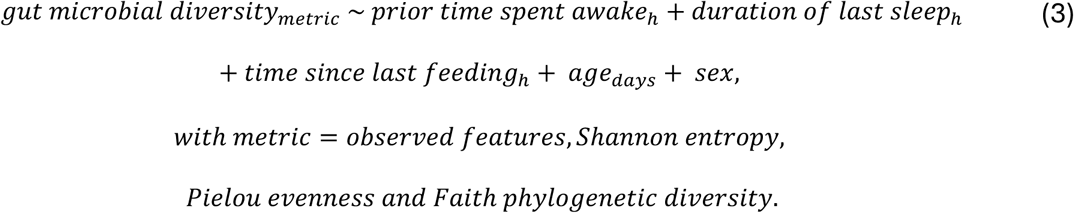

### Modulation of infant sleep by the gut microbiota

To investigate the modulation of infant sleep by the gut microbiota, we examined the associations between gut microbial diversity and temporal volatility and the maturation of infant sleep patterns, including rhythmicity and quality.

#### Sleep rhythmicity: CFI

Two LMMs were created adjusting for the infants’ repeated measures, as well as parenting style (BCQ score), infant behavioral developmental stage (ASQ score), age, and sex, following the equations:

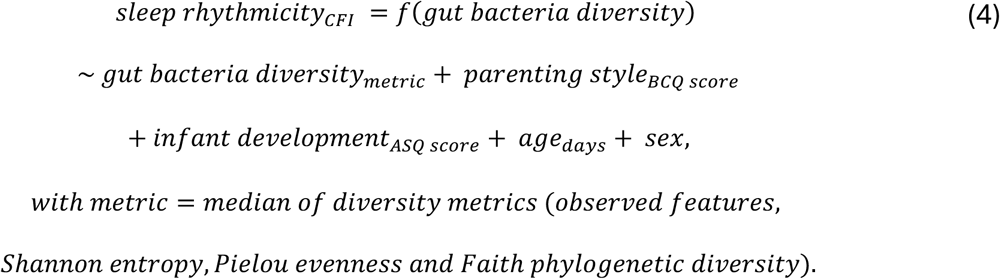

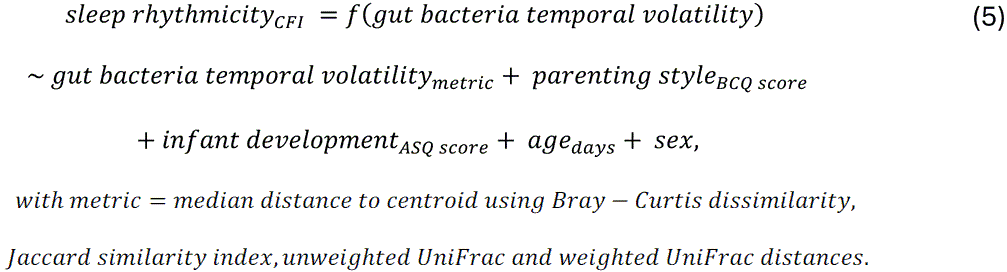

#### Sleep quality: babySQUID

Two LMMs were then created adjusting for the infants’ repeated measures, as well as parenting style (BCQ score), infant behavioral developmental stage (ASQ score), age, and sex, following the equations:

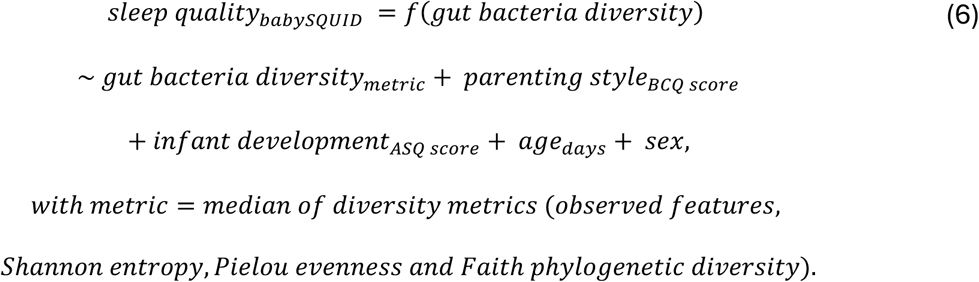

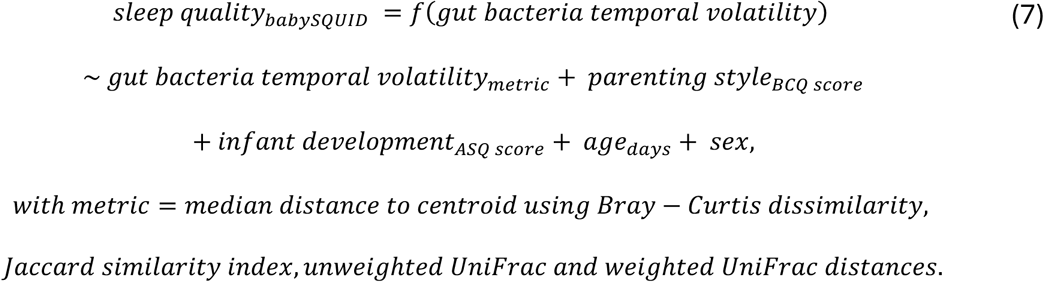

To explore the effect of microbial composition on sleep quality, random forest classifiers were trained to predict the babySQUID score binarized at the median. This approach complements the LMMs, which assume linear relationships, by capturing non-linear complex (high-dimensional) interactions between the gut microbiota and sleep patterns. Input features included (1) the ASV relative abundance table, or (2) its k-merization (Bokulich, 2025), selecting the relative abundance of the top features based on their frequency or TF-IDF score. K-mers length (i.e., n-gram size) was set to 16 as done in previous work (Bokulich, 2025). Models were evaluated using repeated 80/20 train-test splits via different cross-validation methods from Scikit-learn (Pedregosa et al., 2011). Accuracy and weighted F1 (harmonic mean of precision and recall) scores were averaged across the splits.

### Exploratory analysis on gut melatonin

As an exploratory analysis, we tested the association between the abundance of gut melatonin in the samples (in pg/g of stool) and the respective time since the last bowel movement (i.e., the time since the last stool sample), as well as the immediate prior time spent awake (sleep history) and the time since the infant was last fed. A LMM was performed on the samples, adjusting for infants’ repeated measures, age and sex:

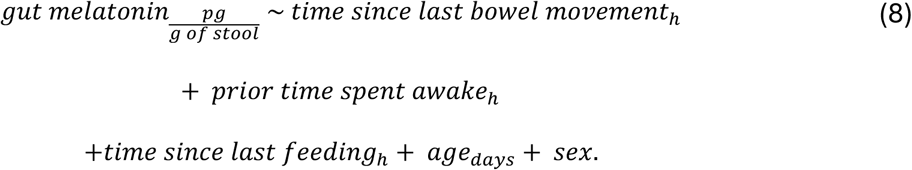

## Results

### Study cohort and microbiota composition

In total, 163 stool samples from 20 infants were included (two samples had missing collection dates and times and 24 samples had a sequencing depth below 3,035 reads, leading to their exclusion; Suppl. Fig. 1). Most infants (n=11) were monitored at two ages, while five infants were assessed at all ages and four infants only had data at one age (2 months: n=11; 4 months: n=15; 6 months: n=15) (Fig. 1, Table 1, Suppl. Fig. 2-3). The gut microbiota of the infant cohort included 426 features from 140 genera, from 10 phyla with *Firmicutes* and *Proteobacteria* being the dominant phyla (Suppl. Fig. 4), with some inter-individual variability (Suppl. Fig. 5). The detailed distribution of the scores from the parent-rated questionnaires on infant sleep (BISQ), parenting style (BCQ), and behavioral development (ASQ) are included in Table 1 and Suppl. Fig. 6-8.

**Table 1.**
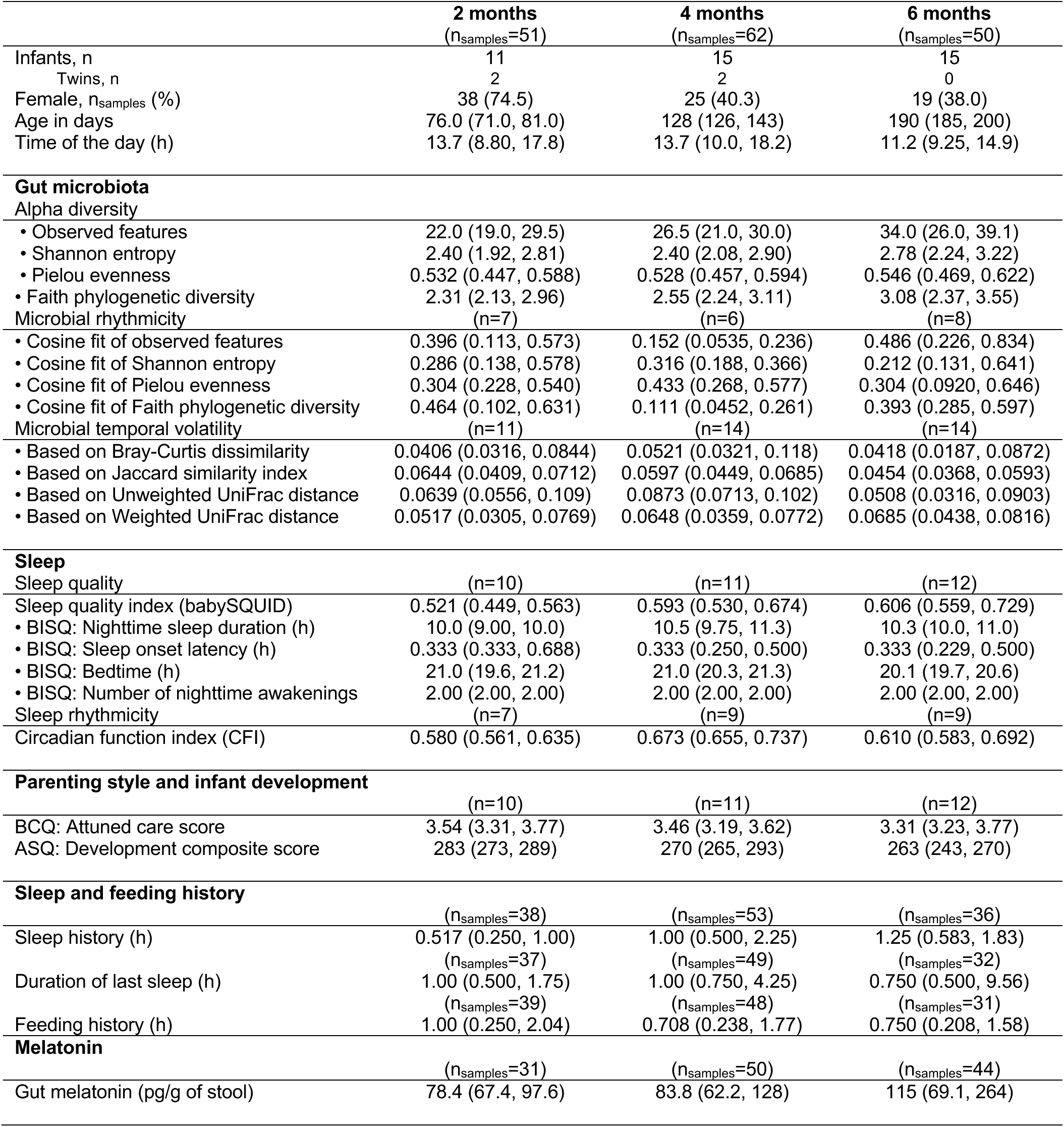
Infants’ samples characteristics. Characteristics of the 163 included samples. Unless otherwise indicated, values represent median (interquartile range).

|  | 2 months<br>(n<sub>samples</sub>=51) | 4 months<br>(n<sub>samples</sub>=62) | 6 months<br>(n<sub>samples</sub>=50) |
| --- | --- | --- | --- |
| Infants, n | 11 | 15 | 15 |
| Twins, n | 2 | 2 | 0 |
| Female, n<sub>samples</sub> (%) | 38 (74.5) | 25 (40.3) | 19 (38.0) |
| Age in days | 76.0 (71.0, 81.0) | 128 (126, 143) | 190 (185, 200) |
| Time of the day (h) | 13.7 (8.80, 17.8) | 13.7 (10.0, 18.2) | 11.2 (9.25, 14.9) |
| **Gut microbiota** |  |  |  |
| Alpha diversity |  |  |  |
| • Observed features | 22.0 (19.0, 29.5) | 26.5 (21.0, 30.0) | 34.0 (26.0, 39.1) |
| • Shannon entropy | 2.40 (1.92, 2.81) | 2.40 (2.08, 2.90) | 2.78 (2.24, 3.22) |
| • Pielou evenness | 0.532 (0.447, 0.588) | 0.528 (0.457, 0.594) | 0.546 (0.469, 0.622) |
| • Faith phylogenetic diversity | 2.31 (2.13, 2.96) | 2.55 (2.24, 3.11) | 3.08 (2.37, 3.55) |
| Microbial rhythmicity |  |  |  |
|  | (n=7) | (n=6) | (n=8) |
| • Cosine fit of observed features | 0.396 (0.113, 0.573) | 0.152 (0.0535, 0.236) | 0.486 (0.226, 0.834) |
| • Cosine fit of Shannon entropy | 0.286 (0.138, 0.578) | 0.316 (0.188, 0.366) | 0.212 (0.131, 0.641) |
| • Cosine fit of Pielou evenness | 0.304 (0.228, 0.540) | 0.433 (0.268, 0.577) | 0.304 (0.0920, 0.646) |
| • Cosine fit of Faith phylogenetic diversity | 0.464 (0.102, 0.631) | 0.111 (0.0452, 0.261) | 0.393 (0.285, 0.597) |
| Microbial temporal volatility |  |  |  |
|  | (n=11) | (n=14) | (n=14) |
| • Based on Bray-Curtis dissimilarity | 0.0406 (0.0316, 0.0844) | 0.0521 (0.0321, 0.118) | 0.0418 (0.0187, 0.0872) |
| • Based on Jaccard similarity index | 0.0644 (0.0409, 0.0712) | 0.0597 (0.0449, 0.0685) | 0.0454 (0.0368, 0.0593) |
| • Based on Unweighted UniFrac distance | 0.0639 (0.0556, 0.109) | 0.0873 (0.0713, 0.102) | 0.0508 (0.0316, 0.0903) |
| • Based on Weighted UniFrac distance | 0.0517 (0.0305, 0.0769) | 0.0648 (0.0359, 0.0772) | 0.0685 (0.0438, 0.0816) |
| **Sleep** |  |  |  |
| Sleep quality |  |  |  |
|  | (n=10) | (n=11) | (n=12) |
| Sleep quality index (babySQUID) | 0.521 (0.449, 0.563) | 0.593 (0.530, 0.674) | 0.606 (0.559, 0.729) |
| • BISQ: Nighttime sleep duration (h) | 10.0 (9.00, 10.0) | 10.5 (9.75, 11.3) | 10.3 (10.0, 11.0) |
| • BISQ: Sleep onset latency (h) | 0.333 (0.333, 0.688) | 0.333 (0.250, 0.500) | 0.333 (0.229, 0.500) |
| • BISQ: Bedtime (h) | 21.0 (19.6, 21.2) | 21.0 (20.3, 21.3) | 20.1 (19.7, 20.6) |
| • BISQ: Number of nighttime awakenings | 2.00 (2.00, 2.00) | 2.00 (2.00, 2.00) | 2.00 (2.00, 2.00) |
| Sleep rhythmicity |  |  |  |
|  | (n=7) | (n=9) | (n=9) |
| Circadian function index (CFI) | 0.580 (0.561, 0.635) | 0.673 (0.655, 0.737) | 0.610 (0.583, 0.692) |
| **Parenting style and infant development** |  |  |  |
|  | (n=10) | (n=11) | (n=12) |
| BCQ: Attuned care score | 3.54 (3.31, 3.77) | 3.46 (3.19, 3.62) | 3.31 (3.23, 3.77) |
| ASQ: Development composite score | 283 (273, 289) | 270 (265, 293) | 263 (243, 270) |
| **Sleep and feeding history** |  |  |  |
|  | (n<sub>samples</sub>=38) | (n<sub>samples</sub>=53) | (n<sub>samples</sub>=36) |
| Sleep history (h) | 0.517 (0.250, 1.00) | 1.00 (0.500, 2.25) | 1.25 (0.583, 1.83) |
|  | (n<sub>samples</sub>=37) | (n<sub>samples</sub>=49) | (n<sub>samples</sub>=32) |
| Duration of last sleep (h) | 1.00 (0.500, 1.75) | 1.00 (0.750, 4.25) | 0.750 (0.500, 9.56) |
|  | (n<sub>samples</sub>=39) | (n<sub>samples</sub>=48) | (n<sub>samples</sub>=31) |
| Feeding history (h) | 1.00 (0.250, 2.04) | 0.708 (0.238, 1.77) | 0.750 (0.208, 1.58) |
| **Melatonin** |  |  |  |
|  | (n<sub>samples</sub>=31) | (n<sub>samples</sub>=50) | (n<sub>samples</sub>=44) |
| Gut melatonin (pg/g of stool) | 78.4 (67.4, 97.6) | 83.8 (62.2, 128) | 115 (69.1, 264) |

### Diurnal patterns: gut microbiota rhythmicity and sleep rhythmicity

Alpha diversity rhythmicity was measured in 21 cases (i.e., instances where infants had at least 4 samples per age) from 16 infants (Suppl. Fig. 9; formula (1)). The estimates of the cosine fits (R^2^ scores) at each age for each diversity metric are displayed in Table 1. For 15 infants the objective sleep rhythmicity information derived from actimetry were available, with a total of 25 CFIs estimated along all ages, based on actimetry recordings with a median duration of 8 days (std= 2.71, range: 3-11 days) (Table 1; formula (2)).

In infants with both datapoints available (14 datapoints from n=12 infants; formula (3)), no significant association was detected between alpha diversity rhythmicity and sleep rhythmicity after adjusting for the infants’ repeated measures, feeding rhythmicity, parenting style, age and sex (p>0.05, Suppl. Fig. 10). Interestingly, even though we observed apparent increases, alpha diversity rhythmicity and sleep rhythmicity were also not significantly associated with age, after adjusting for repeated measures in the infants and the sex (p>0.05, Suppl. Fig. 11-12). However, in all cases there are (insignificant) positive trends of association between age, alpha diversity rhythmicity, and sleep rhythmicity (Fig. 2).

**Fig. 2.**
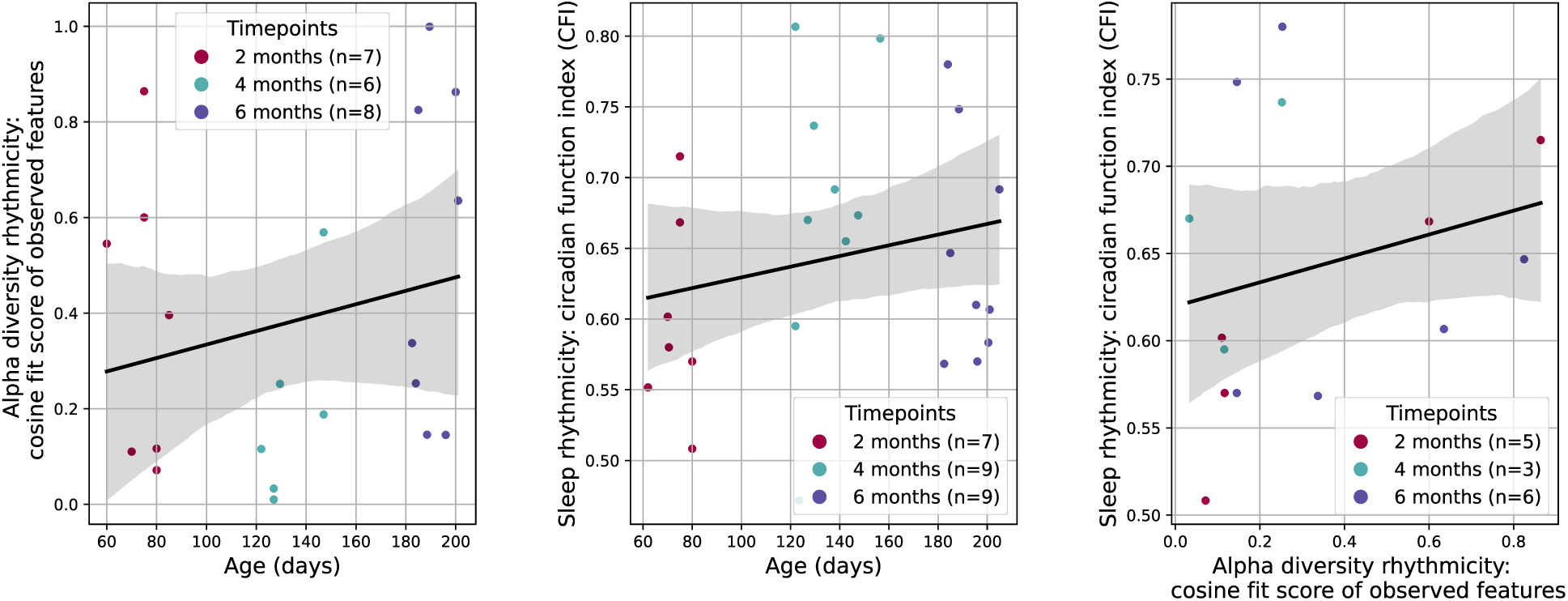
Alpha diversity rhythmicity and sleep rhythmicity in infants. No significant associations between alpha diversity rhythmicity and age, sleep rhythmicity and age, nor sleep rhythmicity and alpha diversity rhythmicity (p>0.05). Regression lines (95% confidence intervals).

### Microbial temporal volatility and age

An indicator for bacterial community instability – gut microbiota temporal volatility – was estimated in all 20 infants at each age (39 datapoints; 2 cases excluded (only 1 sample)) (Table 1, Suppl. Fig. 13). Older infants had lower microbiome temporal volatility, especially when measured using the unweighted UniFrac distance, a measure of the fraction of unique branch length, with β_age_ = –2.81*10^-4^, SE = 1.35*10^-4^, p<0.05 (Fig. 3**A**, Suppl. Fig. 13-14). In other words, as infants grow up, their microbiota become less volatile over 48h (i.e., more stable). Female infants also had less volatile gut microbiomes, when adjusting for the infants’ repeated measures and the age (Temporal volatility (Bray-Curtis dissimilarity): β_male_ = 0.0450, SE = 0.0170, p<0.01, Fig. 3; Suppl. Fig. 14).

**Fig. 3.**
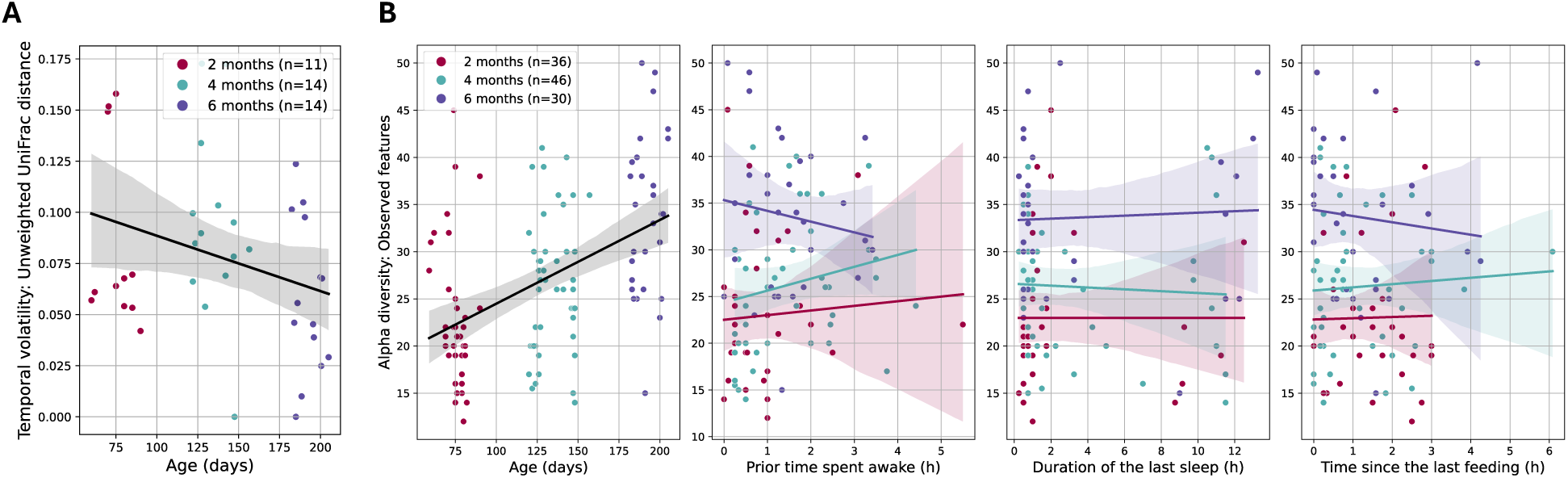
Gut microbiota temporal volatility and alpha diversity in infants. **A** Regression lines (95% confidence intervals) of microbial temporal volatility based on the unweighted UniFrac distance along age (p<0.05). **B** Regression lines (95% confidence intervals) of features abundance along age (p<0.001) and directly preceding sleep and feeding history.

### Effects of sleep and feeding history on the gut microbiota

All sleep and feeding history data were available for 112 samples (Table 1, Suppl. Fig. 15; formula (4)). Fig. 3**B** shows the associations between alpha diversity (measured by feature abundance) and the previous sleep and feeding history in all infants. All four gut microbiota alpha diversity metrics were positively associated with age (observed features: β_age_ = 0.101, SE = 0.0172, p<0.001, Suppl. Fig. 16-17; Fig. 3**B**), when adjusting for the infants’ repeated measures and the sex. However, alpha diversity was associated with neither sleep nor feeding history (p>0.05, Suppl. Fig. 16; Fig. 3**B**). Faith phylogenetic diversity had limited evidence of an association with the infants’ sex, with more species biodiversity present in females (β_male_ = –0.418, SE = 0.237, p<0.1, Suppl. Fig. 16).

### Modulation of infant sleep by the gut microbiota

#### Sleep rhythmicity: CFI

Next, we investigated the association between bacterial alpha diversity and sleep rhythmicity (n=25; formula (5)). We observed higher feature abundance (β_observed features_ = 6.87*10^-3^, SE = 2.54*10^-3^, p<0.01) and microbial biodiversity (β_Faith’s phylogenetic diversity_= 0.0721, SE = 0.0337, p<0.05) in infants with increased CFI (Fig. 4), when controlling for confounders. No associations were detected for Shannon entropy and Pielou evenness (p>0.05, Suppl. Fig. 18). In contrast, there was no association between gut microbiota temporal volatility (bacterial community instability in an infant) and sleep rhythmicity (CFI) (n=24; p>0.05, Suppl. Fig. 19; formula (6)).

**Fig. 4.**
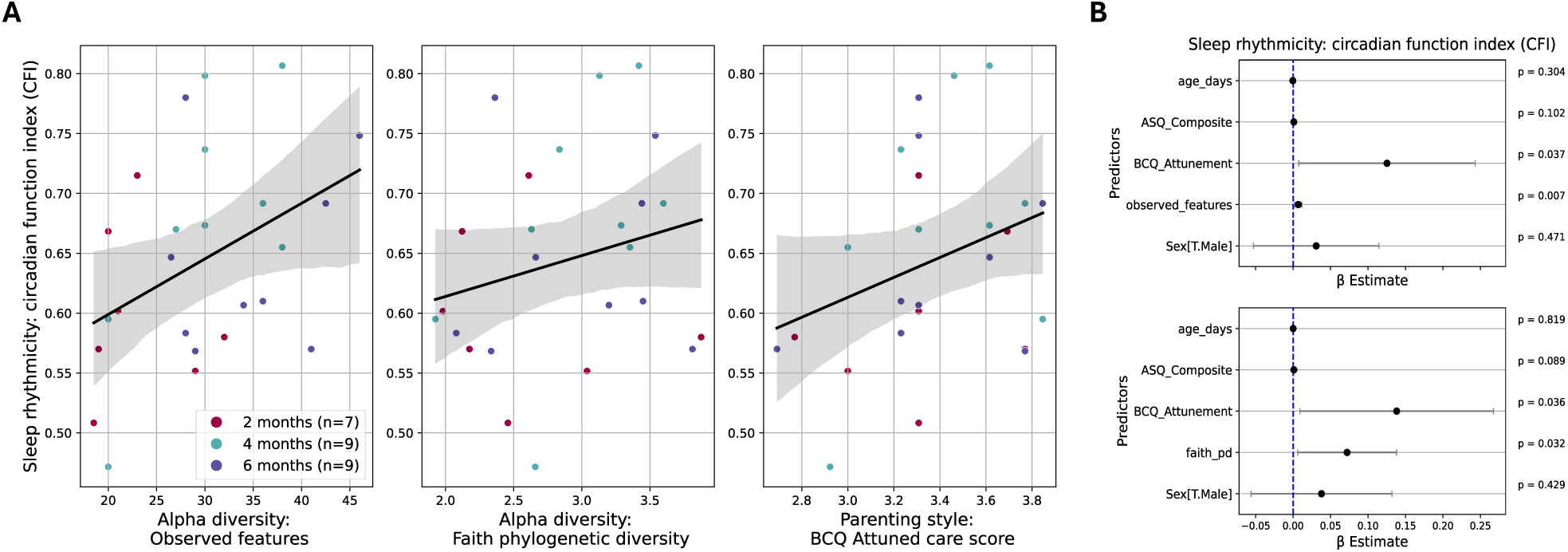
Sleep rhythmicity (CFI) in infants. **A** Alpha diversity measured by features abundance (observed features) and features biodiversity (Faith phylogenetic diversity). Parenting style measured by the attunement score of the BCQ. Regression lines (95% confidence intervals). **B** Coefficient plots. Significant associations between features abundance and biodiversity, and the parenting style, with sleep rhythmicity (p<0.05). Dots are the coefficients, and error bars represent the 95% confidence intervals.

Further, the relevance of behavioral and parenting context was revealed, such that in several models (alpha diversity: observed features and Faith phylogenetic diversity; temporal volatility: Bray-Curtis dissimilarity) there was limited evidence of positive associations between infants with a more attuned care (observed features: β_BCQ_= 0.125, SE = 0.0601, p<0.05), as well as a higher development score (Faith’s phylogenetic diversity: β_ASQ_= 1.03*10^-3^, SE = 6.05*10^-4^, p<0.1), and stronger sleep rhythmicity (Fig. 4 and Suppl. Fig. 19), after adjusting for confounders. The results suggest that, on top of greater alpha diversity, a more attuned caring style and a higher development score are also associated with advanced sleep rhythmicity.

#### Sleep quality: babySQUID

As subjective sleep parameter, the babySQUID was estimated in 33 cases from 18 infants (7, 7, and 4 infants had data at respectively 1, 2 or 3 ages) (Table 1). These are all infants with ≥ 2 infant sleep BISQ scores (excluding two entry errors: reported sleep duration of 16h at 2 months and sleep onset latency of 15h at 2 months). The babySQUID was positively associated with age in all models (Pielou evenness: β_age_ = 1.04*10^-3^, SE = 3.79*10^-4^, p<0.01, Fig. 5), when adjusting for the infants’ repeated measures and the sex (formula (7)). However, there were no significant associations between alpha diversity and sleep quality (babySQUID) in the infants (p>0.05, Fig. 5 and Suppl. Fig. 20). Still, there were positive trends of association between microbial diversity and sleep quality (Fig. 5; β_Shannon entropy_= 0.0577, SE = 0.0320, p<0.1; β_Pielou evenness_= 0.318, SE = 0.192, p<0.1) after adjusting for potential confounders (infants’ repeated measures, parenting style, development scores, age, and sex). No effect of gut microbiota temporal volatility on sleep quality existed in the infants (p>0.05, Suppl. Fig. 21) indicating the association was primarily driven by age (formula (8)). When k-merized, microbial composition predicted the babySQUID (sleep quality) with an accuracy of 63.3% (±8.46%) (Suppl. Table 1), indicating performance above random chance (weakly moderate performance) and supporting the potential utility of microbial profiles as predictive markers of sleep quality.

**Fig. 5.**
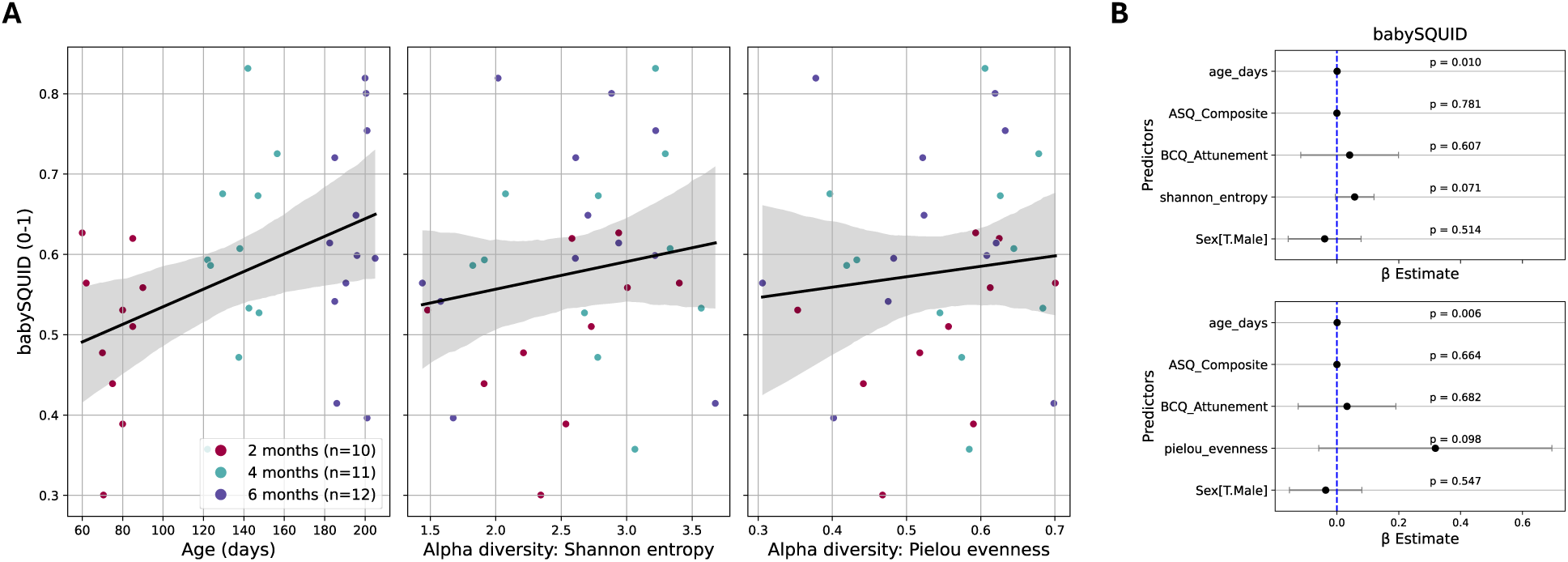
Sleep quality (babySQUID) in infants. **A** Alpha diversity measured by features abundance and evenness (Shannon entropy) and features evenness (Pielou evenness). Regression lines (95% confidence intervals). **B** Coefficient plots. Significant associations between age and the babySQUID (p<0.05); (insignificant) positive trends of association between Shannon entropy and Pielou evenness, and the babySQUID (p<0.1). Dots are the coefficients and error bars represent the 95% confidence intervals.

### Exploratory analysis of gut melatonin

For samples with measured gut melatonin content, the time since the last bowel movement was estimated; note that one melatonin concentration was excluded as an extreme outlier (>8 standard deviations above the mean) (n_samples_=76 from 17 infants; Table 1, Suppl. Fig. 22). Gut melatonin concentration increased with age (β_age_ = 0.638, SE = 0.162, p<0.001, Fig 6; formula (9)) when adjusting for infants’ repeated measures and the sex. No significant association occurred between the time since the last bowel movement, the sleep history, nor the feeding history, and melatonin abundance in stool (p>0.05, Suppl. Fig. 23). Still, in older infants, melatonin showed negative trends of associations along the time since the last bowel movement and sleep history at 6 months, but positive trends for the feeding history (Fig 6).

**Fig. 6.**
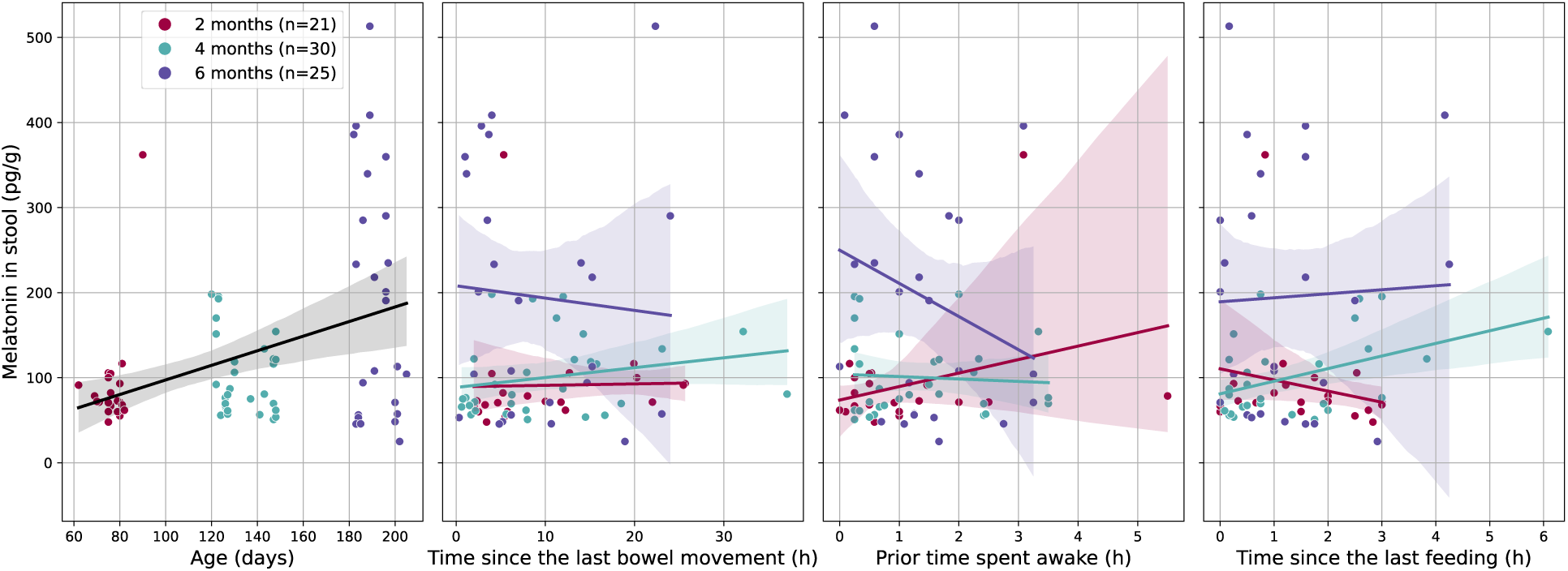
Factors affecting gut melatonin in infants. Regression lines (95% confidence intervals) of melatonin abundance along age (p<0. 001), time since the last bowel movement, the prior time spent awake of the infant, and the time since the last feeding.

## Discussion

To examine associations between the gut microbiota and sleep rhythms in a vulnerable developmental period of infancy, this study combined two methodological strengths: (i) longitudinal within-subject assessments at 2, 4, and 6 months of age combined with (ii) high-frequency sampling and continuous monitoring over a 48-hour period at each age. We used 16S rRNA gene sequencing to examine the gut microbiota, including diversity, rhythmicity and temporal volatility patterns, and combined objective actimetry-based measures of sleep rhythmicity with subjective infant sleep metrics (24-hour diaries, questionnaires) alongside stool melatonin measurement. As hypothesized, findings revealed rhythmicity patterns within a daily timescale, *i.e.*, diurnality, in the gut microbiota in some infants, yet these were not associated with sleep rhythmicity, as quantified by a proxy of circadian rhythm (CFI). As expected, older infants had lower microbiome temporal volatility, indicating less variability in the microbiota of an infant (across 48h). While microbial diversity was also influenced by age, as expected, it was surprisingly not affected by the infant’s directly preceding sleep or feeding history. Also as expected, infants with higher alpha diversity had stronger sleep rhythmicity (CFI) and yet only limited evidence for increased sleep quality (babySQUID). Bacterial temporal volatility was only weakly associated with sleep patterns. These findings suggest that the gut microbiome and sleep patterns are linked in early infancy, with several associations observed across age.

Exploring rhythmicity patterns, as expected, some infants exhibited diurnal rhythms in their gut microbiota, as measured by a cosine fit of alpha diversity. Surprisingly, in contrast to previous studies (Heppner et al., 2024; Mühlematter et al., 2025), no associations between microbiota rhythmicity and infant circadian function index (CFI, sleep rhythmicity) were detected. Still, we report promising positive trends of association between alpha diversity rhythmicity and sleep rhythmicity. Findings also suggested that rhythmicity patterns – in both the microbiota and sleep – may be modulated by age, with positive (though statistically insignificant) trends. The observed increase in microbial rhythmicity with age aligns with previous evidence showing a rise in rhythmic bacterial taxa during the first year of life (Heppner et al., 2024). This also supports the use of the CFI as a reliable proxy for circadian maturation as early as infancy (Mühlematter et al., 2025).

Consistent with expectations, gut bacterial community stability increased with age, indicated by decreased temporal volatility. The effect was particularly pronounced when temporal volatility was assessed using metrics that capture community composition independently of the abundance: the unweighted UniFrac distance and Jaccard similarity index. This suggests that, as infants grow older, multiple samples collected from the same infant at a given age become more similar to one another. Microbiota temporal volatility is a novel marker introduced in this study. We consider it a promising complement to traditional diversity measures, providing additional insights into microbiome stability during early development.

Healthy infants are typically characterized by a low gut bacterial diversity at birth (dominance of *Bifidobacterium*, especially in breast-fed infants) (Lawson et al., 2020), with increasing richness and diversity over time, reaching an adult-like diversity at 14-24 months of age (Stewart et al., 2018; Wernroth et al., 2022). In alignment with this, age was the strongest predictor of gut microbiota diversity in our data (Heppner et al., 2024; Robertson et al., 2019). The influence of immediate context by sleep or feeding history was limited in this study, such that the prior time spent awake, last sleep duration and last feeding time had no effect on the microbiota diversity. An explanation could be that in infants the diet type possibly outweighs the effect of fasting (i.e., time since last feeding) on microbial diversity, in contrast to adults (Zeb et al., 2020). Breast milk directly transmits bacteria and HMOs from the mother to the infant (Selma-Royo et al., 2024), leading to major differences in gut microbiome composition of (exclusively) breast fed and formula fed infants (Robertson et al., 2019). The relationship between sleep recency and duration with alpha diversity in infants is emerging and further studies should be done to properly understand the results (Zimmermann et al., 2025). While studies in adults and animal models have already demonstrated associations between gut microbiota composition and sleep patterns, there is further need for well-controlled interventional trials in infancy that can test causal pathways in the gut-brain-sleep axis.

Investigating the links between diurnal rhythmicity patterns in the gut microbiota and infant sleep rhythmicity, higher gut bacterial diversities were associated with stronger sleep rhythmicity (as measured by the CFI) in the infants. Notably, we observed higher gut bacterial abundance and microbial biodiversity in infants with a stronger sleep rhythmicity, when controlling for confounders. This expands on and supports the results from previous studies (Al-Andoli et al., 2025; Mühlematter et al., 2025; Schoch et al., 2022), adding a focus on the association between infant gut microbial diversity and sleep rhythmicity. However, this pattern contrasts with findings in other domains of early development, where lower microbial diversity has been associated with more favorable outcomes (Alderete et al., 2021; Moroishi et al., 2022). This highlights the need for more nuanced modeling approaches that account for non-linear maturation trajectories, age-specific effects, and potentially even group-level differences. Surprisingly, temporal volatility in the gut microbiota (over 48-hours) was not associated with differences in sleep rhythmicity. By quantifying short-term fluctuations in microbiota composition, this approach aimed to provide a high-resolution view of microbial dynamics in relation to behavioral rhythms such as sleep, yet no significant association was observed in this sample.

We found no evidence of an association between microbiota diversity and sleep quality, as defined by the babySQUID (combining infant sleep variables from the parent-reported BISQ). While studies have investigated the effect of sleep patterns (e.g. evening bedtime) on microbial diversity at a later age (Olson et al., 2024), none have replicated our analysis. Gut microbiota temporal volatility was also not a marker for better sleep quality in this study and further research should be performed to investigate the associations between gut temporal volatility and sleep quality. Random forest classifiers predicted sleep quality (babySQUID) with moderate accuracy, achieving best performance and lowest variability when using the top 20 k-mers. As k-mers capture pseudo-phylogenetic sequence-level patterns beyond taxonomic resolution (Bokulich, 2025), this indicates that a targeted subset of microbial features – rather than reductive metrics such as alpha diversity – may carry the most predictive power of sleep quality in the infants. This suggests that gut microbiota composition, rather than diversity alone, could be a marker for infant sleep quality.

An exploration of factors affecting gut melatonin in the samples revealed that its abundance increased with age in the infants, in line with previous findings (Al-Andoli et al., 2025). Accumulation may result from either the infants’ own system or from maternal sources. Endogenous production via the pineal gland is more likely in older infants, although accumulated melatonin from maternal milk may also contribute (Häusler et al., 2024). Still, in older infants, gut melatonin levels tended to decrease with increasing time since the last bowel movement and with longer periods spent awake. We previously observed (insignificant) decreases in species diversity with increasing time since the last sleep and feeding at 6 months. This suggests that bacterial metabolic activity could be elevated shortly after these events and then gradually declines over time.

Investigating other potential factors influencing infant sleep, we detected that in addition to greater alpha diversity, a more attuned caring style and a higher development composite score were also associated with more robust sleep rhythmicity. This suggests that the infants’ environment is also important in shaping their sleep. Furthermore, this aligns with the link between sleep and brain development, e.g. linking behavioral and neurophysiological patterns of sleep with neuronal connectivity (Schoch et al., 2022, 2024). Interestingly, after adjusting for age, microbiome temporal variability – and to some extent, diversity – was greater in females, suggesting that biological sex, along with feeding and environmental factors, might influence microbiota development (Robertson et al., 2019). Associations between microbiota development and biological sex have not been reported or found insignificant in most prior studies of term infants (Stewart et al., 2018), but have been reported as significant in a longitudinal study of pre-term infants (Chen et al., 2021) and could warrant further investigation.

This study represents a pioneering effort in understanding the associations between the gut microbiota, sleep and feeding patterns in infants, by means of high-resolution, continuous sampling and integrative analysis across objective and subjective infant data. The strengths of this pilot study include continuous sampling (n_samples_=163) with good sequencing resolution (minimum sequencing depth of 3,035 reads per sample). Specifically, high-frequency sampling (median=4 samples per infant per age (std= 2.45, min=1, max=11)) is unique to this investigation, standing in contrast to cross-sectional or sparse longitudinal designs (one sample per infant per age) assessed in prior studies. Multiple samples per age allow for a unique look at diurnal rhythms of the gut microbiota with reduced variability (providing low variability in the results), while accounting for individual differences. Moreover, this study used objective and subjective dimensions of sleep: actimetry, 24-h-diaries, and the BISQ. Objective sleep data came from continuous measures (wearable tracking) averaged over several days (CFI) as a proxy for circadian rhythm in the infant. Finally, all analyses were rigorously controlled for potential confounders (age, sex, feeding history, parenting style and infant behavioral developmental stage), adding some robustness in the results.

To further clarify the relationship between the microbiome and sleep in infancy, larger studies with longer follow-up periods are needed to capture even broader individual developmental trajectories, microbiome maturation patterns, and potential causal directions in the association between gut composition and sleep regulation. In this study, estimating gut microbiota diversity rhythmicity was only possible in 16 infants at a specific age (i.e., age where at least 4 samples were collected within a 48-h period). The low sample size is a result of the multi-measure and demanding continuous monitoring protocol required but reduces statistical power to detect potential rhythmicity associations (type 2 error). Moreover, this study used objective sleep data as a proxy for circadian rhythm in the infant, unlike previous approaches (Heppner et al., 2024). The rhythmicity analysis should be replicated in a study with a larger sample size to confirm our promising findings. Analyses on specific taxa (e.g., *Bifidobacteria*, major metabolizers of HMOs) could also be investigated and bring some explanations on the mechanistic pathways involved. Future randomized control studies should investigate causal mechanisms, test these relationships in more diverse or at-risk populations, and explore the long-term implications for infant (neuro)development (Zimmermann et al., 2025). Moreover, the infant sleep quality index (babySQUID) included information on sleep onset latency and bedtime, not linearly associated with better sleep quality. Indeed, a fast sleep onset latency could be indicative of an overall lack of sleep, while earlier bedtimes may not generally reflect infants with better sleep quality. In addition, we expect infant sleep to be influenced by the real-life context or cultural/social circumstances including factors that were not monitored in this study. Finally, the microbiome-sleep relationship appears to be bidirectional, which could confound interpretations of the present findings and highlight the need for longitudinal or interventional studies to disentangle the directionality of this relationship (Sen et al., 2021).

This study highlights the intricate relationship between gut microbiota rhythmicity, diversity and temporal volatility, with infant sleep rhythmicity and quality. It also integrates information on feeding patterns, parenting care style, behavior development, and gut melatonin. With both a diurnal and longitudinal resolution, this study provides a foundation for future research, offering insights into the potential role of the gut microbiota in shaping sleep rhythmicity and sleep consolidation in infants. Looking ahead, these findings open up promising avenues for targeted interventions – ranging from dietary strategies to microbiota-based therapies – that could support healthy sleep and (neuro)development in infants.

## Acknowledgments

We thank the participants and parents for their dedicated help in this study. The authors also thank Natalie Bakir and Nadja Steinke for their help with assessments and data handling.

## Author contributions

Conceptualization, S.K. and N.A.B.; methodology, F.K., C.M., A.A., S.K., and N.A.B.; validation, S.K. and N.A.B.; formal analysis, F.K. and C.M.; investigation, C.M., D.F., S.P., and P.Z.; resources, S.K. and N.A.B.; data curation, F.K. and C.M.; writing – original draft, F.K., S.K., and N.A.B.; writing – review and editing, F.K., C.M., A.A., D.F., S.P., P.Z., S.K., and N.A.B.; visualization, F.K.; supervision, S.K. and N.A.B.; project administration, S.K. and N.A.B.; funding acquisition, S.K. and N.A.B.

## Declaration of generative AI and AI-assisted technologies in the writing process

During the writing process, after completing the initial draft, the authors used ChatGPT (OpenAI) to improve the writing style and correct spelling and grammar in the manuscript. Following the use of this tool, the authors carefully reviewed and edited the content as necessary and take full responsibility for the final published research article.

## Resource availability

### Materials availability

This study did not generate new unique reagents.

### Data and code availability

- Newly generated sequencing data (16S rRNA) from stool samples in this study will be made publicly available in the NCBI Sequence Read Archive (SRA) as of the date of publication.
- All original code will be deposited on a repository and publicly available at as of the date of publication.
- Any additional information required to reanalyze the data reported in this paper is available from the corresponding authors upon request.

