## Supplementary Materials for "The gut microbiota and sleep in infants: a focus on diurnal rhythmicity patterns"

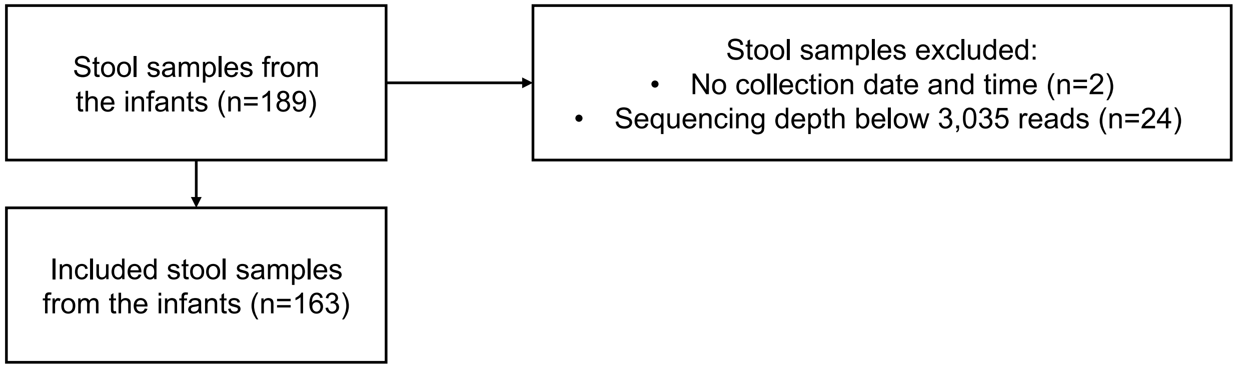


Suppl. Fig. 1. **Flow chart of the inclusion and exclusion process of samples.**

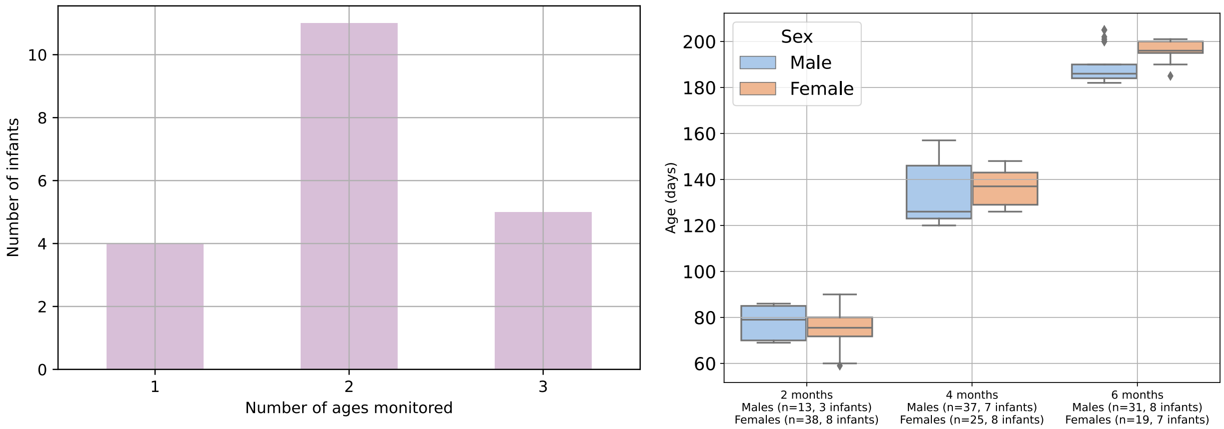


Suppl. Fig. 2. **(left) Distribution of infants by assessment ages. (right) Infants' age and sex distribution across ages.**

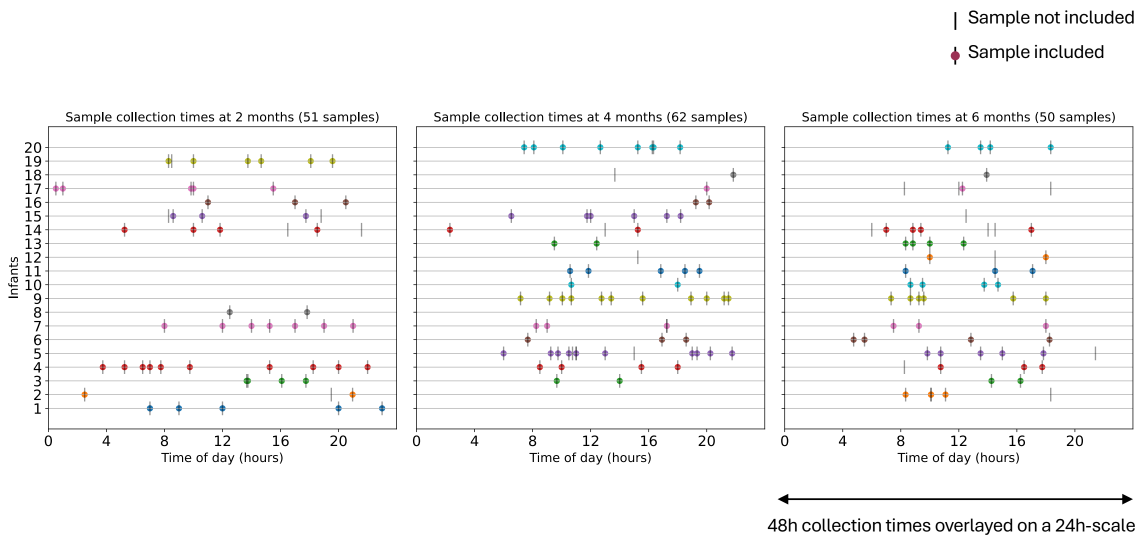


Suppl. Fig. 3. **Stool samples collection times across all ages and infants.**

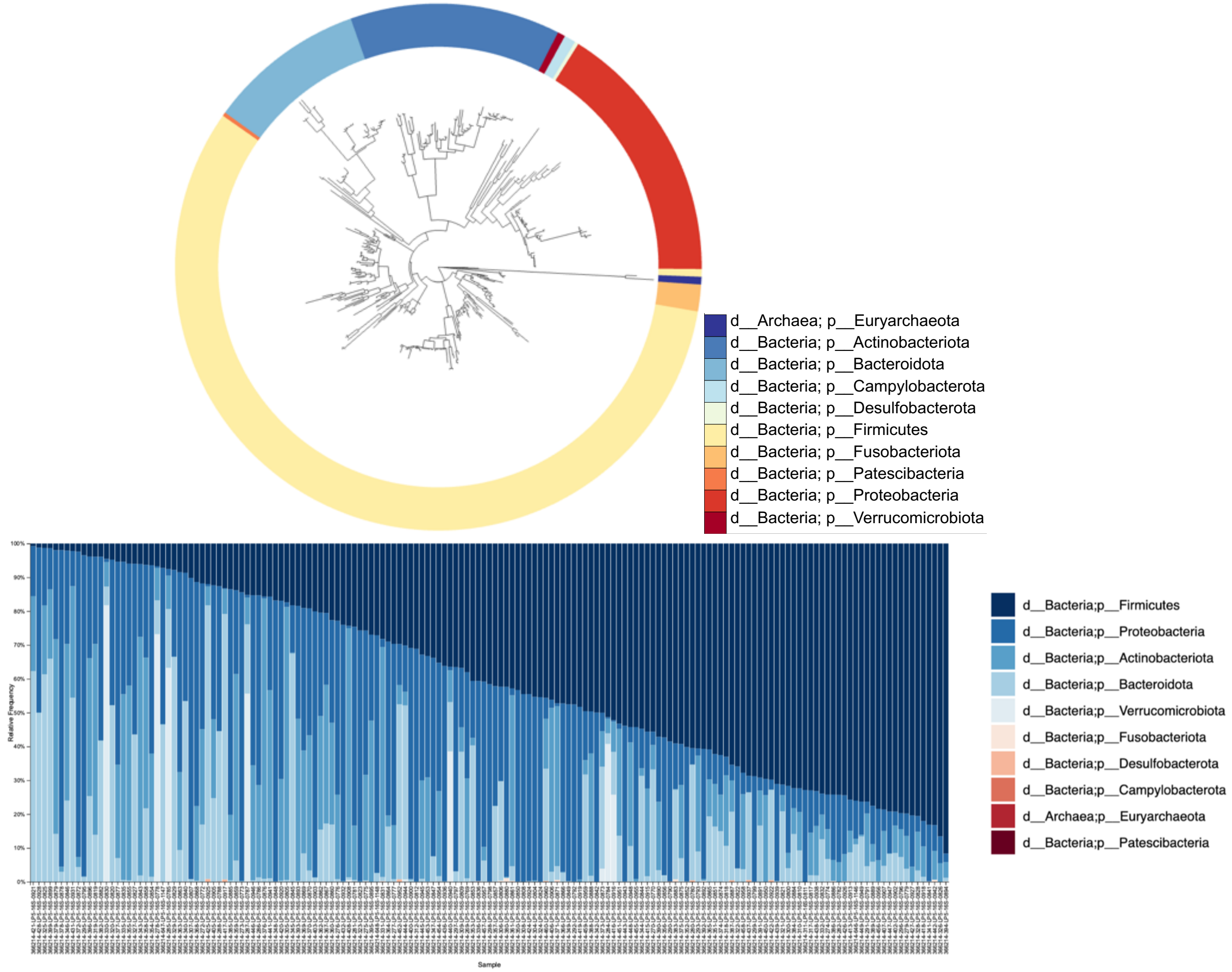


Suppl. Fig. 4. **Gut microbiota composition**. (top) Phylogenetic tree and (bottom) stacked bar plots of the phyla present in the samples.

**A**

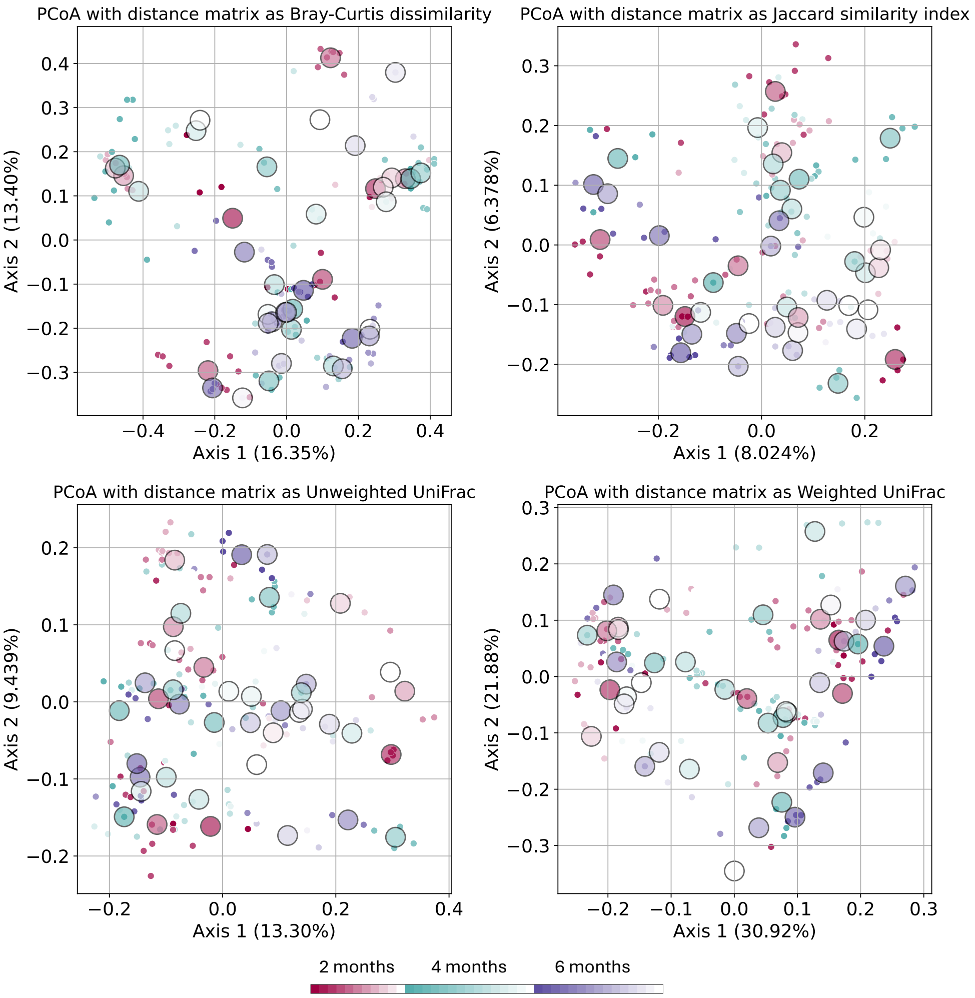


**B**

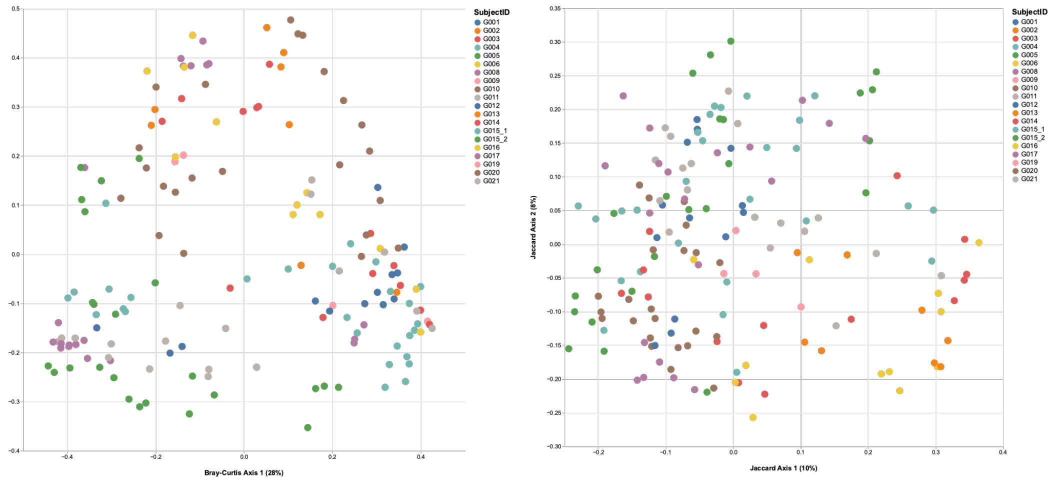


Suppl. Fig. 5. **Gut microbiota beta diversity (between-samples (dis)similarity)**. Each color is an infant. **A** Features measured using ASVs. **B** Features measured using k-mers.

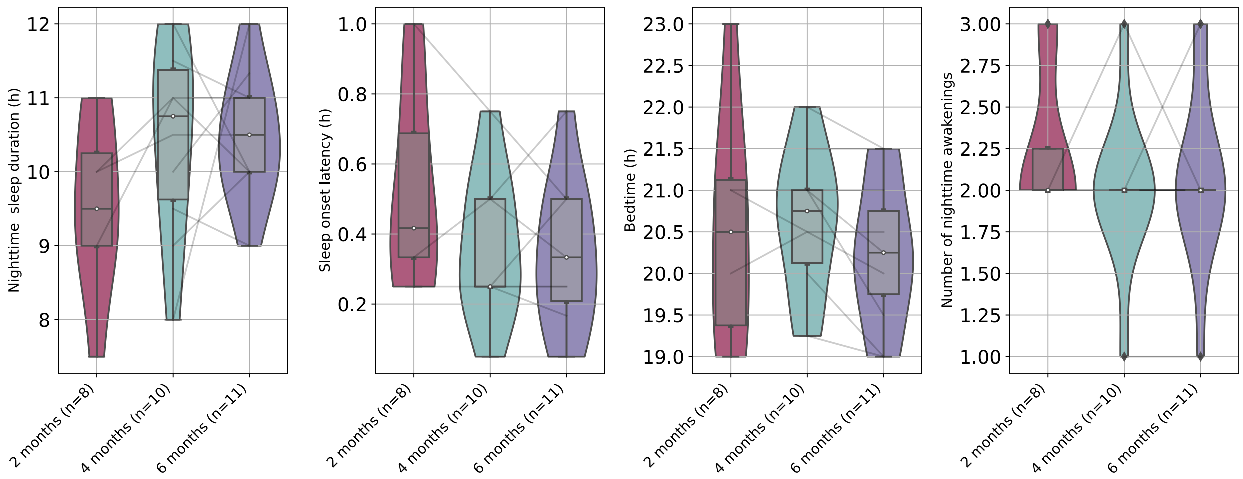


Suppl. Fig. 6. **Infant sleep BISQ variables across all ages.** Sleep duration, sleep onset latency, bedtime and frequency of waking in the night variables.

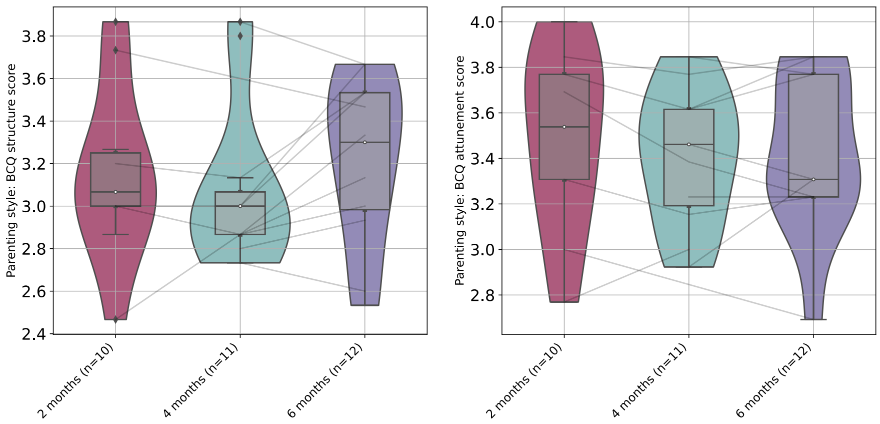


Suppl. Fig. 7. **BCQ variables across all ages.** Structured and attuned care scores.

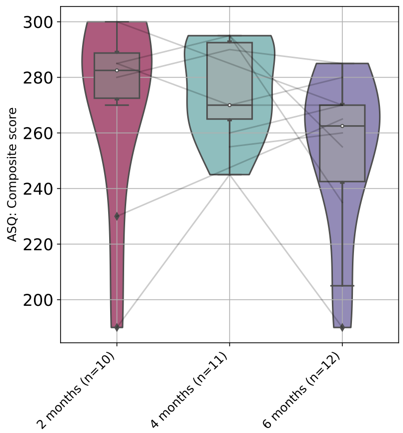


Suppl. Fig. 8. **ASQ Composite score across all ages.** Represents communication, fine motor, gross motor, personal-social and problem solving scores.

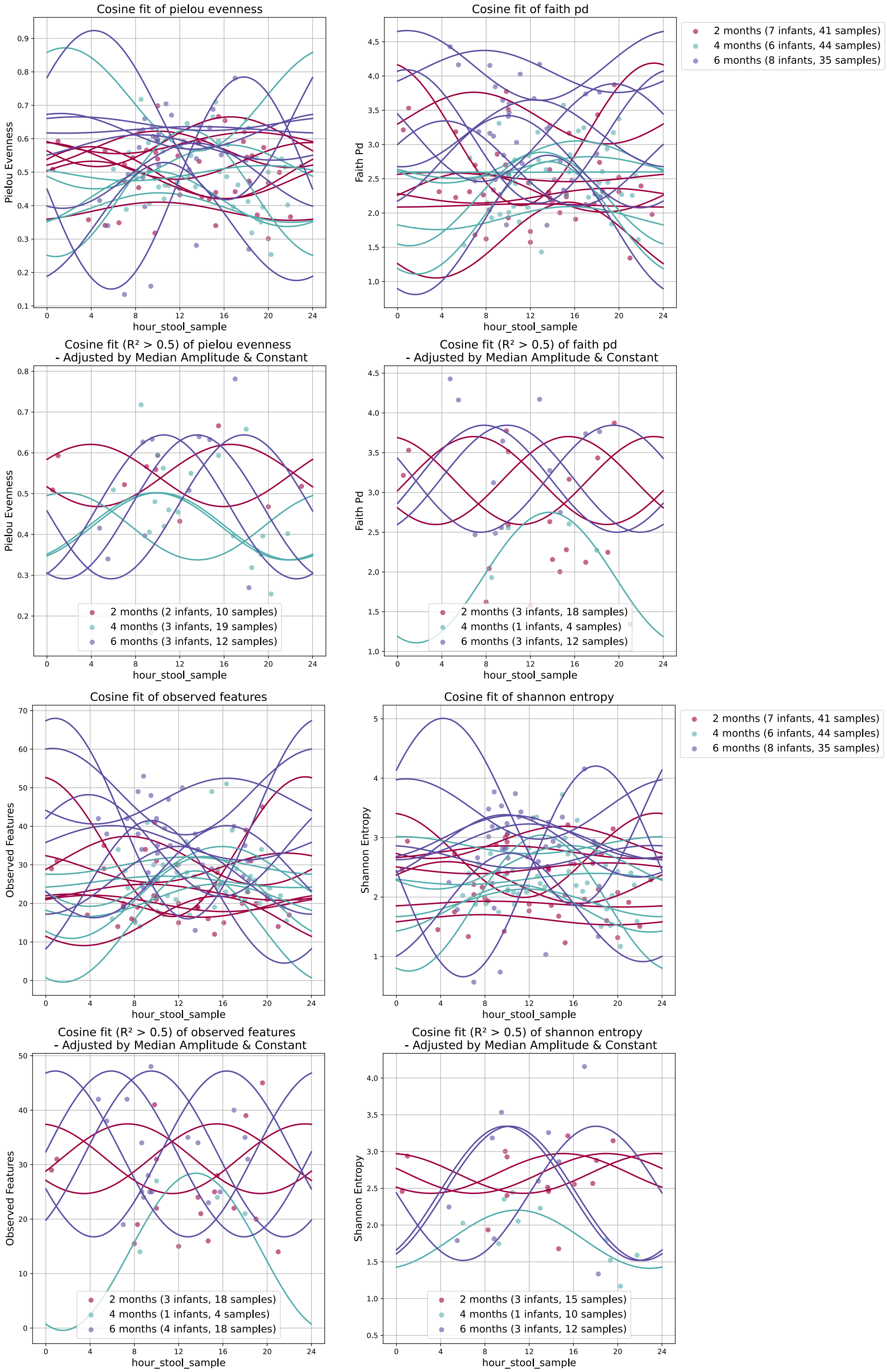


Suppl. Fig. 9 **Cosine fitting of alpha diversity metrics across age.** Cosine functions are plotted for the infants having at least 4 samples per age.

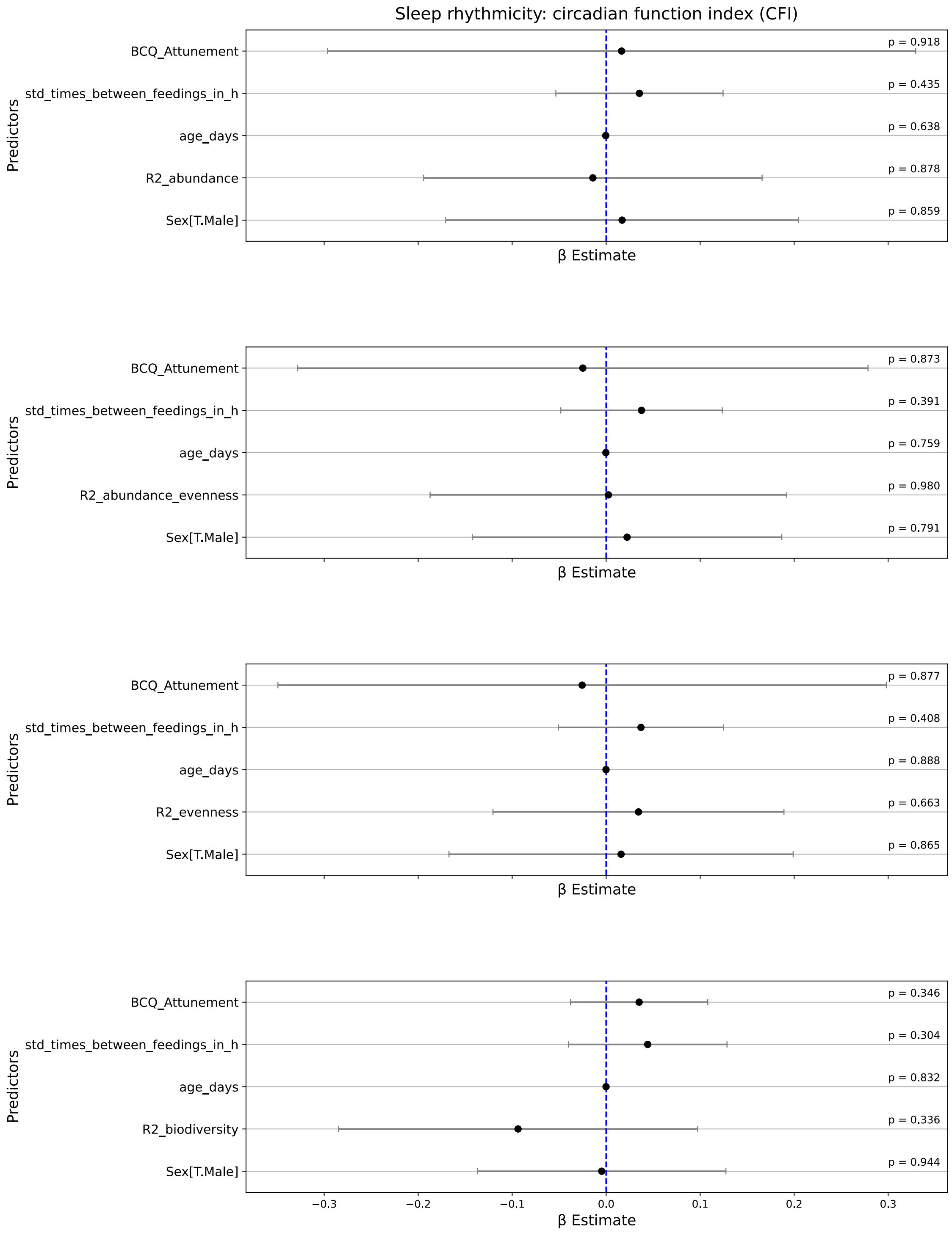


Suppl. Fig. 10. **Coefficient plots showing alpha diversity rhythmicity effects on sleep rhythmicity (n=14).** No significant associations (p>0.05). R2_abundance, R2_abundance_evenness, R2_evenness and R2_biodiversity respectively correspond to the $R^2$ values (measures of cosine fit) of alpha diversity metrics observed features, Shannon entropy, Pielou evenness, and Faith's phylogenetic diversity. Dots are the coefficients and error bars represent the 95% confidence intervals.

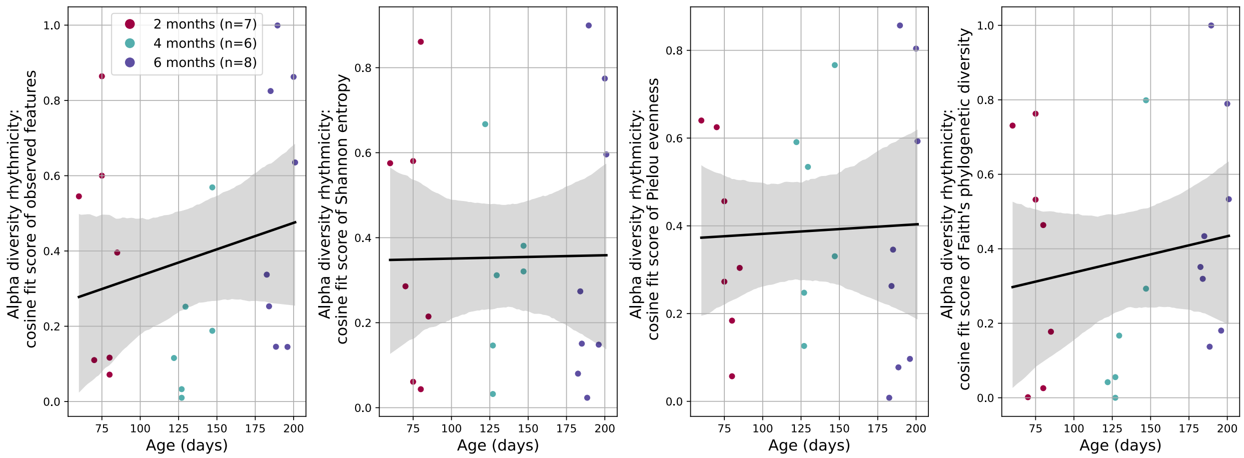


Suppl. Fig. 11. **Alpha diversity rhythmicity and age**. Regression lines (95% confidence intervals) of rhythmicity metrics along age. Alpha diversity rhythmicity measured by the cosine fit ($R^2$ score) of observed features, Shannon entropy, Pielou evenness and Faith's phylogenetic diversity (p>0.05). Only infants having at least 4 samples per age are included.

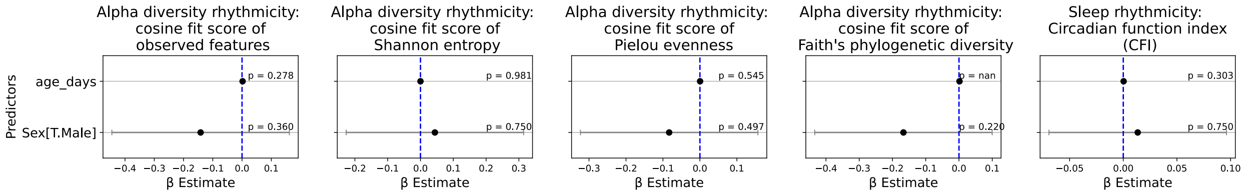


Suppl. Fig. 12. **Coefficient plots showing age and sex effects on alpha diversity rhythmicity (n=21) and sleep rhythmicity (n=25).** No significant associations (p>0.05). The p-value for age_days of the fourth model (Faith Phylogenetic diversity) is reported as NaN most probably due to insufficient within-subject variation: 11 infants had only one datapoint, which limits the model's ability to estimate the within-subject effect of the variables. Dots are the coefficients and error bars represent the 95% confidence intervals.

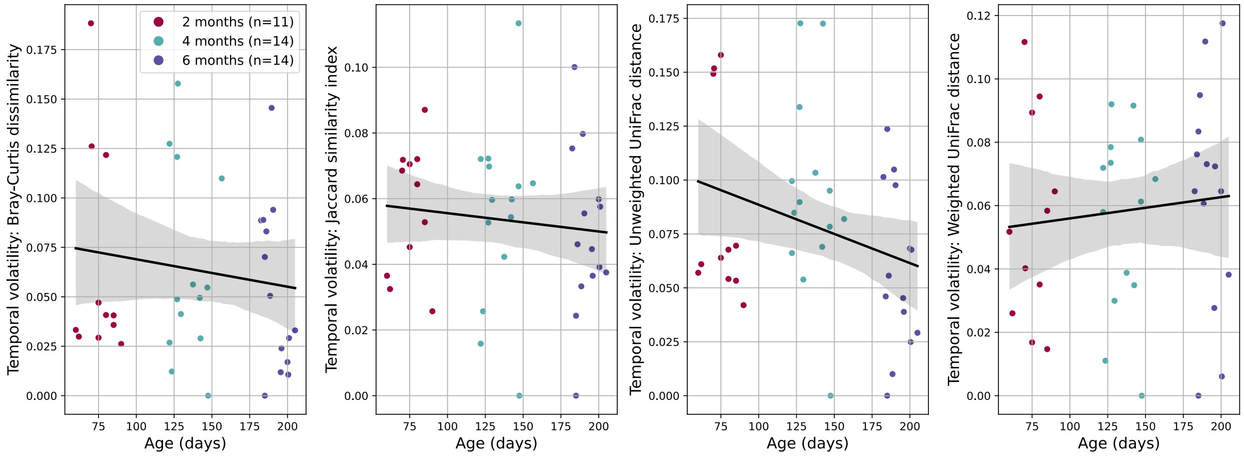


Suppl. Fig. 13. **The gut microbiota temporal volatility and age**. Regression lines (95% confidence intervals) of temporal volatility metrics along age; significant association between the unweighted UniFrac distance and age (p<0.05). Only cases with at least 2 samples per age are included.

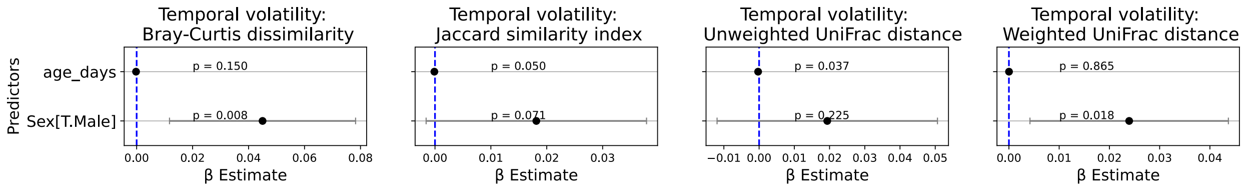


Suppl. Fig. 14. **Coefficient plots showing age and sex effects on microbial temporal volatility.** Significant associations for both age and sex in some models (p<0.05). Dots are the coefficients and error bars represent the 95% confidence intervals.

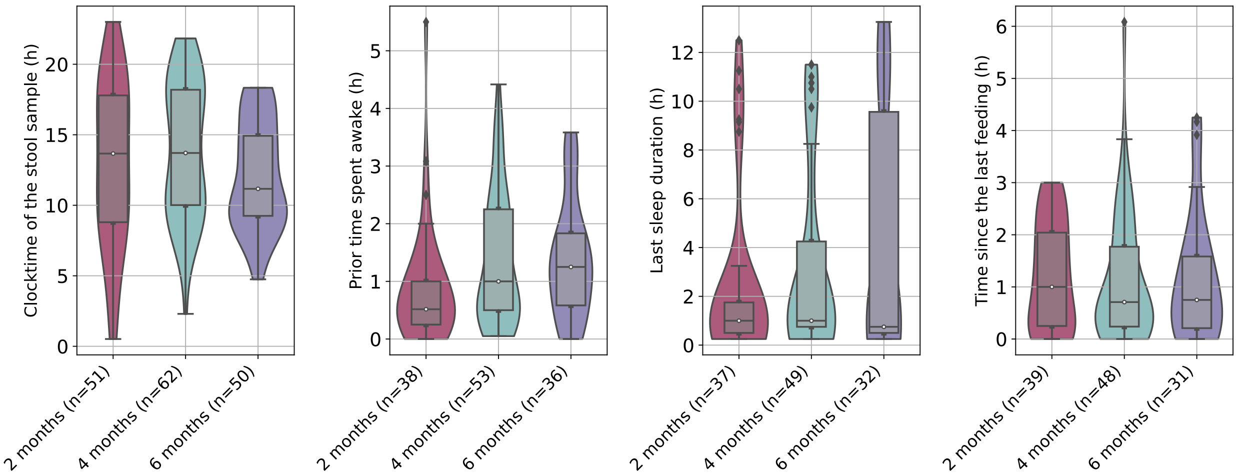


Suppl. Fig. 15. **Clock times of stool samples and sleep and feeding history at each stool sample time**. Visualization is limited to samples with directly preceding feeding and sleeping information.

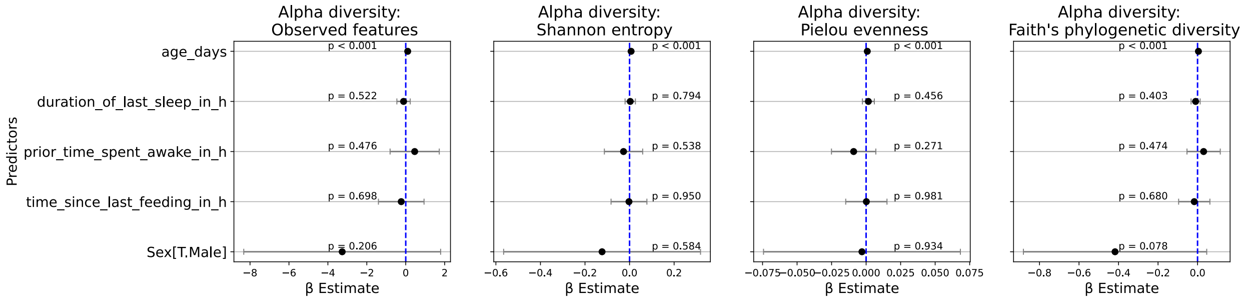


Suppl. Fig. 16. **Coefficient plots showing the sleep and feeding history effects on alpha diversity.** Significant associations between all alpha diversity metrics and age (p<0.001). Limited evidence of an association between features biodiversity (Faith phylogenetic diversity) and sex (p<0.1). Dots are the coefficients and error bars represent the 95% confidence intervals.

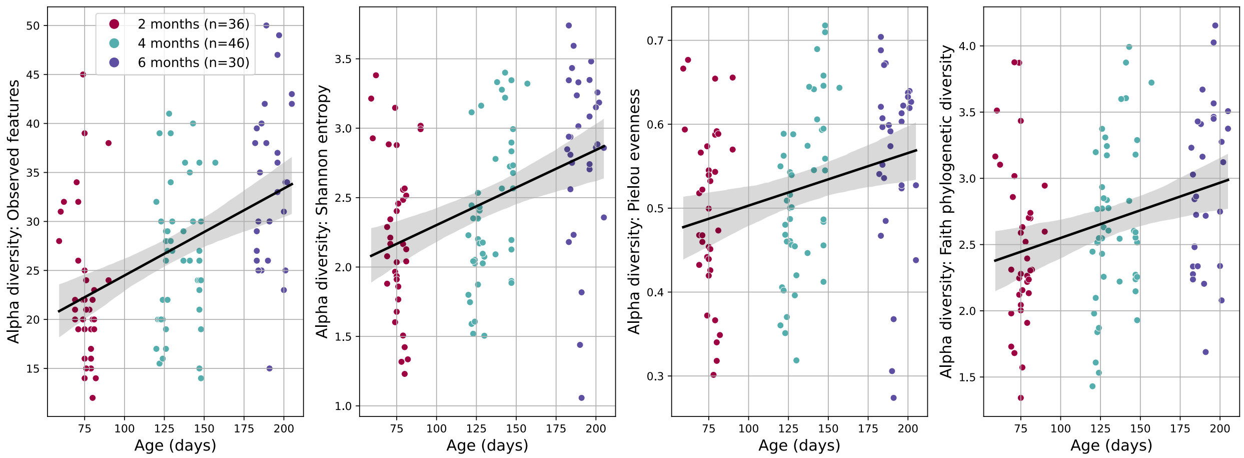


Suppl. Fig. 17. **Alpha diversity and age**. Regression lines (95% confidence intervals) of alpha diversity metrics (observed features, Shannon entropy, Pielou evenness and Faith phylogenetic diversity) along age in the samples ($p<0.001$).

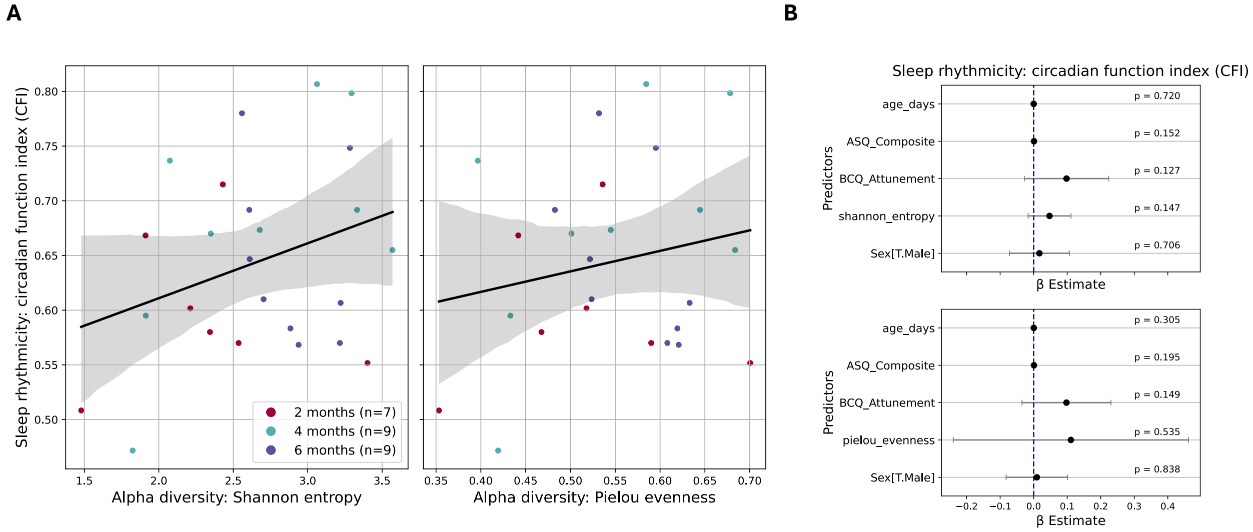


Suppl. Fig. 18. **Sleep rhythmicity (CFI) and alpha diversity**. **A** Regression lines (95% confidence intervals). Alpha diversity measured by features abundances and evenness (Shannon entropy) and features evenness (Pielou evenness). **B** Coefficient plots. No significant associations ($p>0.05$). Dots are the coefficients and error bars represent the 95% confidence intervals.

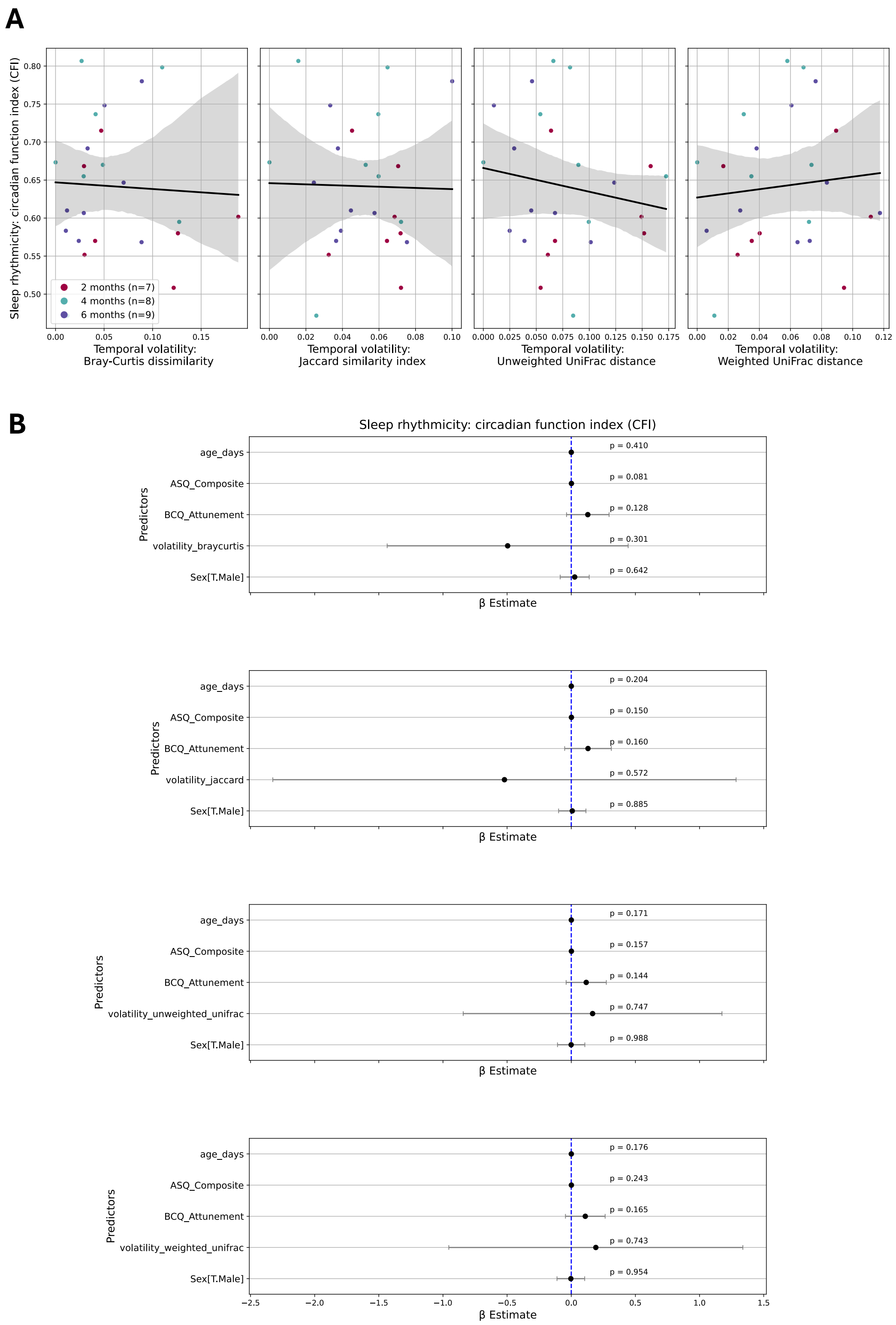


Suppl. Fig. 19. **Sleep rhythmicity and gut microbiota temporal volatility**. **A** Regression lines (95% confidence intervals). Gut microbiota temporal volatility measured by Bray-Curtis dissimilarity, Jaccard similarity index, Unweighted UniFrac and Weighted UniFrac distances. **B** Coefficient plots. No significant associations (p>0.05). Dots are the coefficients and error bars represent the 95% confidence intervals.

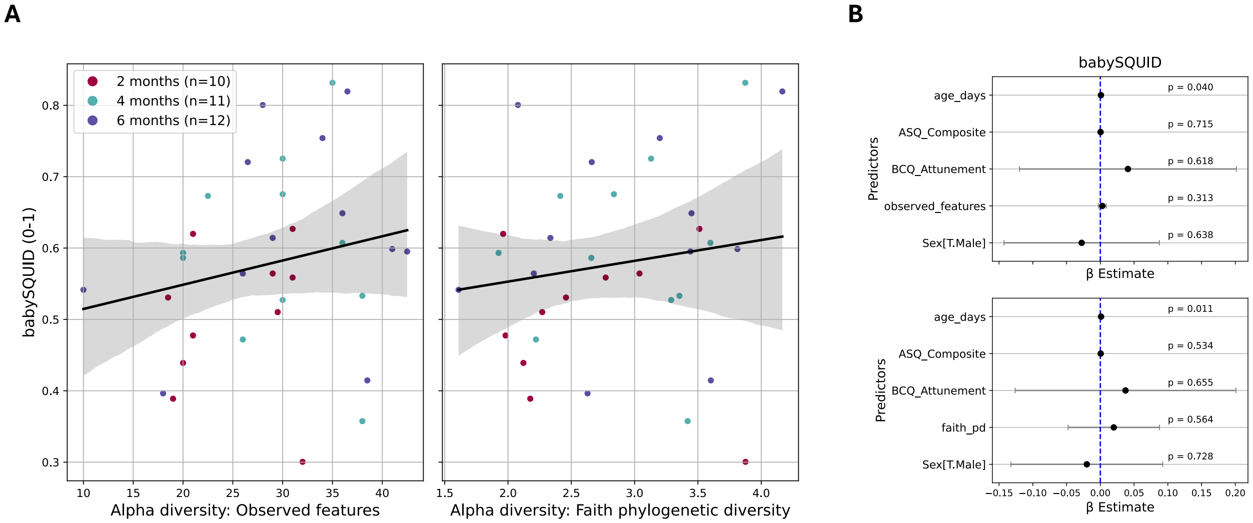


Suppl. Fig. 20. **Sleep quality (babySQUID) and alpha diversity**. **A** Regression lines (95% confidence intervals). Alpha diversity measured by features abundance and biodiversity (Faith phylogenetic diversity). **B** Coefficient plots. Significant associations between age and sleep quality ($p<0.05$). Dots are the coefficients and error bars represent the 95% confidence intervals.

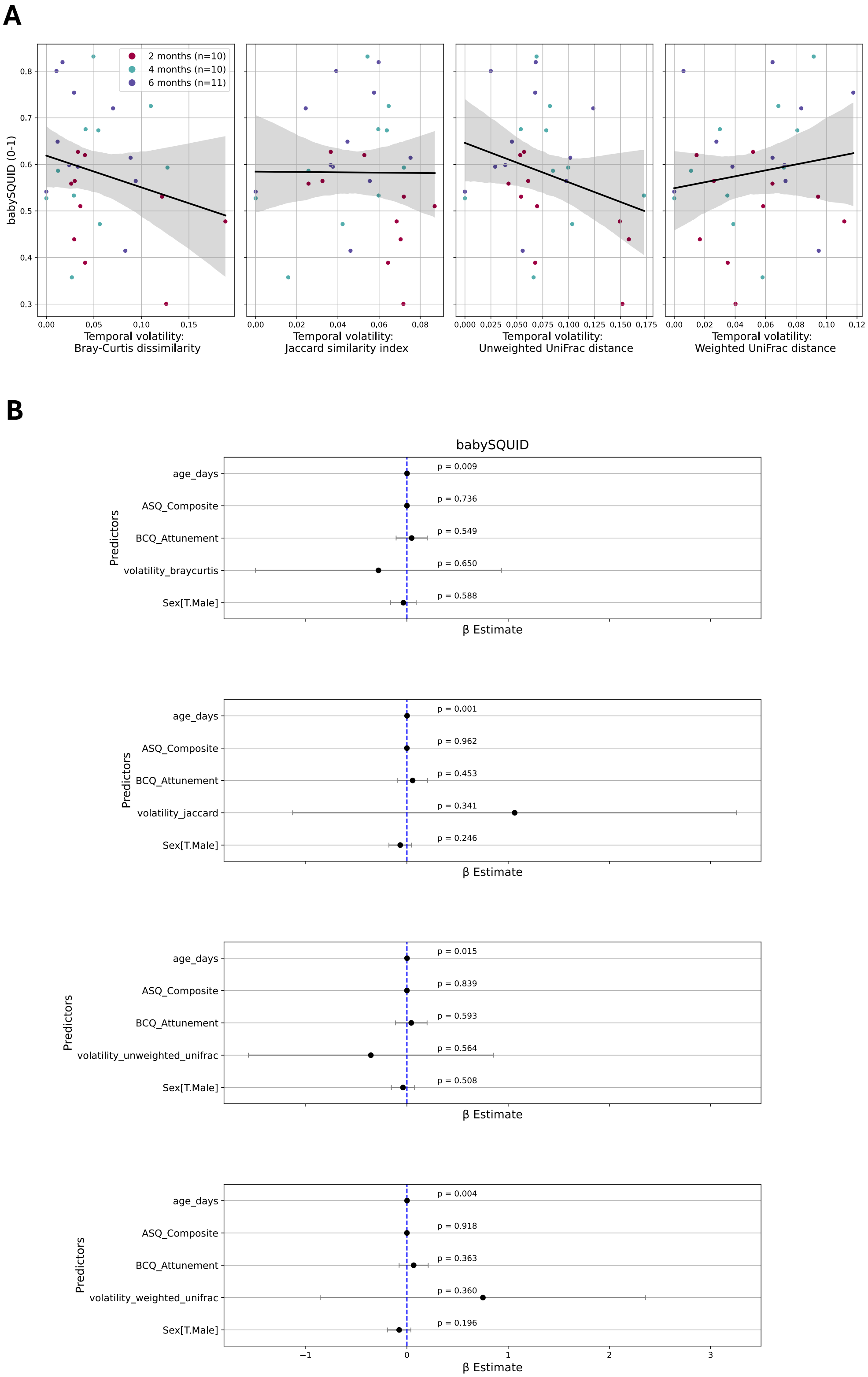


Suppl. Fig. 21. **The babySQUID and gut microbiota temporal volatility. A** Regression lines (95% confidence intervals). Gut microbiota temporal volatility measured by Bray-Curtis dissimilarity, Jaccard similarity index, Unweighted UniFrac and Weighted UniFrac distances. **B** Coefficient plots. Significant associations between age and sleep quality ($p<0.05$). Dots are the coefficients and error bars represent the 95% confidence intervals.

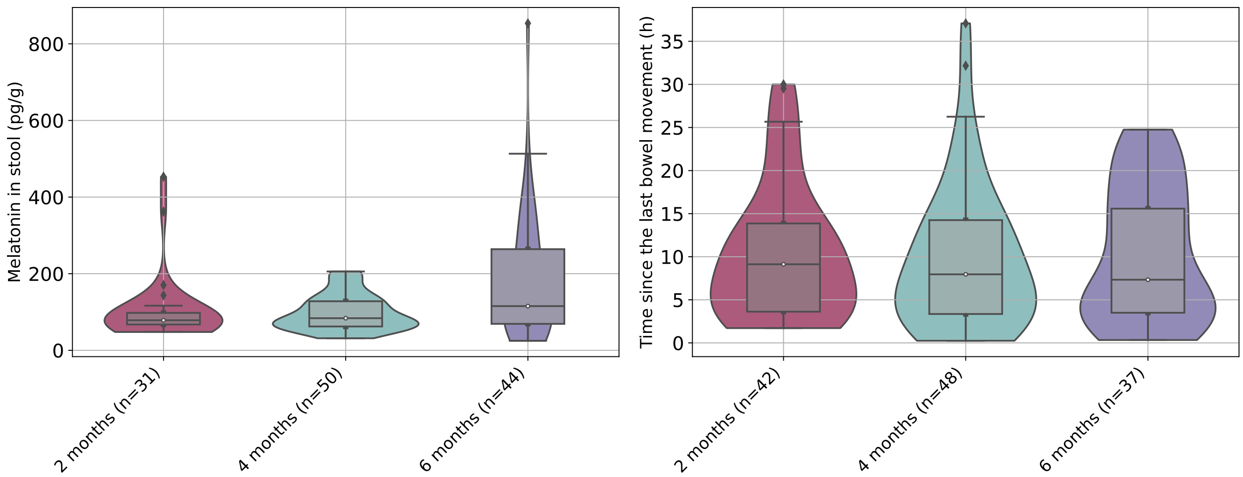


Suppl. Fig. 22. **Gut melatonin and time since the last bowel movement at each stool sample.** Visualization limited to samples with melatonin ($n_{samples}$=125) and previous stool sample ($n_{samples}$=127) information.

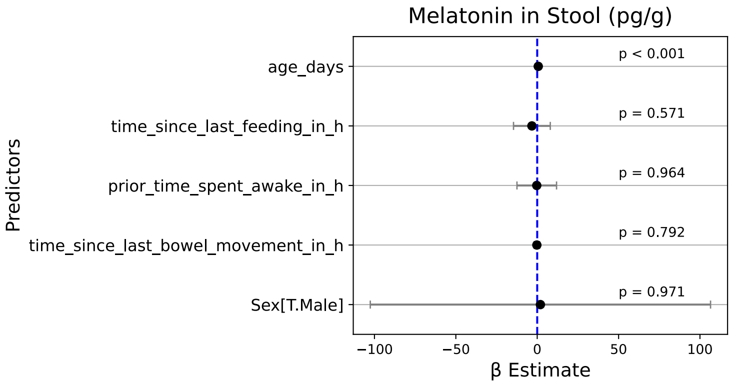


Suppl. Fig. 23. **Coefficient plots showing factors effects on gut melatonin.** Melatonin abundance positively associated with age ($p<0.001$). Dots are the coefficients and error bars represent the 95% confidence intervals.

| Feature Table | Cross-Validation | Mean accuracy (± SD) | Mean weighted F1 Score (± SD) |
|---|---|---|---|
| **ASVs (426 features)** | | | |
| | GroupShuffleSplit | 0.468 ± 0.174 | 0.381 ± 0.225 |
| | GroupKFold | **0.586 ± 0.249** | **0.551 ± 0.277** |
| | LeaveOneGroupOut | 0.593 ± 0.402 | 0.574 ± 0.424 |
| **K-mers (k=10)** | | | |
| | GroupShuffleSplit | 0.408 ± 0.156 | 0.372 ± 0.159 |
| | GroupKFold | 0.424 ± 0.105 | 0.370 ± 0.100 |
| | LeaveOneGroupOut | 0.306 ± 0.417 | 0.298 ± 0.419 |
| **K-mers (k=20)** | | | |
| | GroupShuffleSplit | 0.598 ± 0.042 | 0.595 ± 0.040 |
| | GroupKFold | **0.633 ± 0.085** | **0.609 ± 0.103** |
| | LeaveOneGroupOut | 0.574 ± 0.409 | 0.576 ± 0.416 |
| **K-mers (k=50)** | | | |
| | GroupShuffleSplit | 0.438 ± 0.061 | 0.431 ± 0.082 |
| | GroupKFold | 0.548 ± 0.080 | 0.493 ± 0.137 |
| | LeaveOneGroupOut | 0.491 ± 0.362 | 0.465 ± 0.375 |
| **TF-IDF k-mers (k=10)** | | | |
| | GroupShuffleSplit | 0.416 ± 0.161 | 0.389 ± 0.180 |
| | GroupKFold | 0.510 ± 0.129 | 0.455 ± 0.158 |
| | LeaveOneGroupOut | 0.444 ± 0.412 | 0.419 ± 0.421 |
| **TF-IDF k-mers (k=20)** | | | |
| | GroupShuffleSplit | 0.544 ± 0.079 | 0.536 ± 0.092 |
| | GroupKFold | **0.633 ± 0.085** | **0.609 ± 0.103** |
| | LeaveOneGroupOut | 0.574 ± 0.374 | 0.557 ± 0.389 |
| **TF-IDF k-mers (k=50)** | | | |
| | GroupShuffleSplit | 0.517 ± 0.103 | 0.520 ± 0.102 |
| | GroupKFold | 0.548 ± 0.080 | 0.505 ± 0.141 |
| | LeaveOneGroupOut | 0.546 ± 0.359 | 0.539 ± 0.375 |

Suppl. Table 1. **Results of random forest classifiers**. Classifiers predicting the babySQUID binarized at the median (0 if below median, 1 if > median). Using the relative abundances of ASVs to predict the babySQUID, the model achieved an average accuracy of 58.6% (±24.9%) and a weighted F1 score of 0.551 (±0.277). Predictive performance improved further after k-merizing the feature table (k-mer length = 16) and selecting the top 20 features. This approach increased the accuracy to 63.3% (±8.46%) and the weighted F1 score to 0.609 (±0.103), indicating both enhanced performance and reduced variability (same results obtained when top features selected based on their frequency and TF-IDF score). Cross-validation methods: (1) GroupShuffleSplit (n=5): generates a user-determined number of random test splits, each with a user-determined fraction of unique groups; (2) GroupKFold (n=5): K-fold iterator variant with non-overlapping groups; (3) LeaveOneGroupOut (n=18 infants): leave-one-out approach (i.e., for each of N folds, train on N-1 infants and test on the Nth, where N = number of infants). Accuracy = (number of correct predictions) / (total number of predictions). F1 Score = harmonic mean of precision and recall (precision = proportion of true positives in all predicted positives; recall = proportion of (correctly) predicted positives out of all true positives).
